# Allosteric links between the hydrophilic N-terminus and transmembrane core of human Na^+^/H^+^ antiporter NHA2

**DOI:** 10.1101/2022.04.26.489501

**Authors:** Diego Velázquez, Vojtěch Průša, Gal Masrati, Hana Sychrova, Nir Ben-Tal, Olga Zimmermannova

**Affiliations:** Laboratory of Membrane Transport, Institute of Physiology of the Czech Academy of Sciences, Prague 4, Czech Republic; Department of Biochemistry and Molecular Biology, George S. Wise Faculty of Life Sciences, Tel-Aviv University, Tel-Aviv, Israel

## Abstract

The human Na^+^/H^+^ antiporter NHA2 (*SLC9B2*) transports Na^+^ or Li^+^ across the plasma membrane in exchange for protons, and is implicated in various pathologies. It is a 537 amino acids protein with an 82 residues long hydrophilic cytoplasmic N-terminus followed by a transmembrane part comprising 14 transmembrane helices. We optimized the functional expression of *Hs*NHA2 in the plasma membrane of a salt-sensitive *Saccharomyces cerevisiae* strain and characterized a set of mutated or truncated versions of *Hs*NHA2 in terms of their substrate specificity, transport activity, localization and protein stability. We identified a highly conserved proline 246, located in the core of the protein, as being crucial for ion selectivity. The replacement of P246 with serine or threonine resulted in antiporters with altered substrate specificity and increased resistance to the *Hs*NHA2-specific inhibitor phloretin that were not only highly active at an acidic pH of 4.0 (like the native antiporter), but also at neutral pH. We also experimentally confirmed the importance of a putative salt bridge between E215 and R432 for antiporter function and structural integrity. Truncations of the first 50 - 70 residues of the N-terminus doubled the transport activity of *Hs*NHA2, whilst changes in the charge at positions E47, E56, K57, or K58 decreased the antiporter’s transport activity. Thus, the hydrophilic N-terminal part of the protein appears to allosterically autoinhibit its cation transport. Our data also show this *in vivo* approach to be useful for a rapid screening of SNP’s effect on *Hs*NHA2 activity.

## Introduction

The maintenance of ion homeostasis is crucial for any living cell, as it influences various physiological parameters such as cell size, intracellular pH, and membrane potential. Cation/H^+^ antiporters (CPAs, SLC9 family) are tightly regulated transporters that ensure appropriate intracellular concentrations of monovalent cations in organisms of all kingdoms, from bacteria to mammals (Padan and Landau, 2016). They mediate the exchange of monovalent cations, mainly Na^+^ and K^+^, for one or two protons across the membrane. Human cation/H^+^ antiporters account for 13 isoforms from three subfamilies – NHE (SLC9A), which includes 9 proteins (NHE1-9), NHA (SLC9B) with two members, NHA1 and NHA2, and the SLC9C subfamily with two proteins (SLC9C1 and SLC9C2) of unknown functions (Masrati et al., 2018; Pedersen and Counillon, 2019). Deficiencies in the functions of CPAs are related to a growing list of pathologies, ranging from hypertension to autism spectrum disorders and cancer, which demonstrates their importance for human health (Pedersen and Counillon, 2019).

In this work we focus on a member of the SLC9B subfamily encoded by the NHA2 gene. It is a Na^+^(Li^+^)/H^+^ antiporter, which is expressed in multiple tissues (Fuster et al., 2008), but its physiological functions are relatively poorly defined. Within individual cells, it was shown to be localized to the plasma membrane or intracellularly to endosomes or mitochondria (Battaglino et al., 2008; Fuster et al., 2008). It generally contributes to the regulation of intracellular pH, sodium homeostasis, and cellular volume, and it was found to play crucial roles in various physiological processes depending on the specific tissue where it resides.

All knowledge obtained so far indicates that NHA2 performs multiple functions in the human body and its malfunctioning leads to various pathologies ranging from metabolic to fertility disorders. Moreover, the NHA2 gene is located in a human chromosomal region (4q24) which has been associated with hypertension in numerous linkage studies (Xiang et al., 2007). So far, particular physiological roles of NHA2 have been studied in a range of organisms and tissues. Single RNAi knockdowns of NHA2 or its homologue NHA1 in *Drosophila melanogaster* reduced survival, and their combination was lethal, suggesting their essentiality for life (Chintapalli et al., 2015). In pancreatic β-cells of mice and humans, NHA2 resides in endosomes and is critical for insulin secretion and clathrin-mediated endocytosis (Deisl et al., 2013). The loss of NHA2 also exacerbates obesity- and aging-induced glucose intolerance in mice (Deisl et al., 2016), and a single-nucleotide polymorphism of the NHA2 locus was recently associated with type 2 diabetes in humans (H. M. Liu et al., 2018). In the kidney, NHA2 localizes to distal convoluted tubules, where it is critical for electrolyte (sodium reabsorption) and blood pressure homeostasis (Fuster et al., 2008; Kondapalli et al., 2017). In 2021, Anderegg et al. corroborated the NHA2 role in blood pressure regulation in mice, and identified it as critical for the maintenance of serine-threonine with-no-lysine kinase 4 (WNK4) levels, and thus ultimately for the activity of the Na^+^/Cl^−^ cotransporter NCC in the distal convoluted tubules of the kidney (Anderegg et al., 2021). Increased NHA2 expression also promotes cyst development in an *in vitro* model of polycystic kidney disease (Prasad et al., 2019). In the testis, NHA2 is relevant for sperm motility and male fertility, since the deletion of NHA2 significantly reduced the percentage of motile sperm (of about 30%) and resulted in a decrease in pregnancy rate (of about 25%) in mice (S. R. Chen et al., 2016). In addition, several studies showed NHA2 to be expressed in osteoclasts, where NHA2 depletion significantly inhibited osteoclast differentiation *in vitro* (Battaglino et al., 2008; Ha et al., 2008; S. H. Lee et al., 2008), however its physiological role *in vivo* in this type of cell remains to be elucidated (Hofstetter et al., 2010; Charles et al., 2012).

As for its structure, human NHA2 is a protein 537 amino acids in length and 58 KDa in size. It consists of 14 transmembrane segments (TMS), a hydrophilic N-terminus approximately 80 amino acids long, and a relatively short (27 aa) hydrophilic C-terminus. A two-dimensional representation of *Hs*NHA2 topology is schematically shown in Fig. 1A, and in detail in Fig. S1. Recently, 3D-structures of human NHA2 and its bison homolog were determined using cryo-electron microscopy (Protein Data Bank 7B4M/L; (Matsuoka et al., 2022), respectively). However, the hydrophilic N-terminal and C-terminal parts are missing from both structures. The N- and C-termini are predicted to be unstructured, for example by the recently published AI-based protein structure prediction algorithm AlphaFold2 (Varadi et al., 2022). The importance of the highly unstructured hydrophilic N-terminal part for the functioning of NHA2 has not been studied yet.

**Figure 1:**
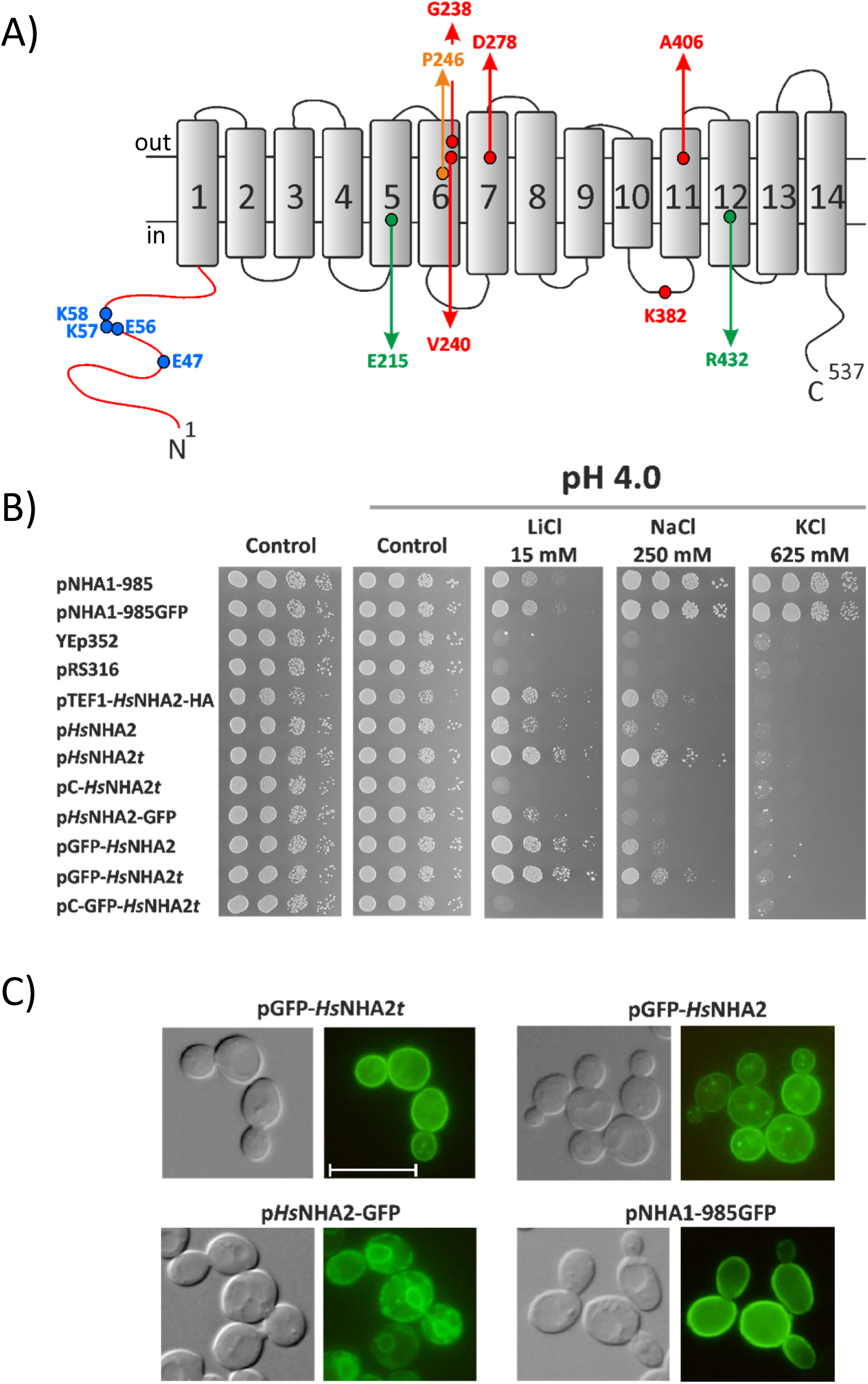
Optimization of conditions for heterologous expression of *Hs*NHA2 in *S. cerevisiae*. A) Schematic representation of transmembrane topology of *Hs*NHA2. The positions of residues studied in this work are highlighted. B) Growth of alkali-metal cation-sensitive BW31 strain containing empty vectors (YEp352, pRS316) or expressing *Sc*Nha1 or *Hs*NHA2 (non-tagged or N- or C-terminally tagged with GFP) from various types of plasmids (Table 1) was compared on YNB-Pro (pH approx. 4.8) or non-buffered YNB-Pro plates (pH 4.0) supplemented with LiCl, NaCl, or KCl as indicated (*t* = *TPS1* terminator, pC = centromeric plasmid). Cells were grown at 30 °C for 2 (control), 6 (LiCl) or 4 (NaCl, KCl) days. C) Nomarski (left) and fluorescence (right) micrographs of BW31 cells expressing either N-terminally GFP-tagged *Hs*NHA2, C-terminally GFP-tagged *Hs*NHA2 or *Sc*Nha1-GFP (expressed from pGFP-*Hs*NHA2*t*, pGFP-*Hs*NHA2, p*Hs*NHA2-GFP or pNHA1-985GFP, respectively). Cells were grown in YNB-Pro (4% glucose) to the early exponential phase. The scale bar corresponds to 10 μm.

*Hs*NHA2 mediates an electroneutral efflux of sodium or lithium cations in exchange for external protons across the membrane (Chintapalli et al., 2015; Uzdavinys et al., 2017). Some studies indicate that, depending on the proton or cation concentrations, it is able to work in both directions and also provides Na^+^/Li^+^ counter transport in kidney-derived MDCK cells (Kondapalli et al., 2012; Kondapalli et al., 2017). Pharmacologically, NHA2 is resistant to amiloride, a typical inhibitor of members of the NHE subfamily, but it is a phloretin-sensitive transport system (Kondapalli et al., 2012; Xiang et al., 2007).

So far, several functional and mutagenesis studies of *Hs*NHA2 have been done upon its expression in the model yeast *Saccharomyces cerevisiae*, in a strain that lacked the three Na^+^ transporters – the Na^+^/H^+^ antiporter Nha1 and Na^+^-ATPases Ena from the plasma membrane and the endosomal Na^+^/H^+^ antiporter Nhx1 (Fuster et al., 2008; Matsuoka et al., 2022; Schushan et al., 2010; Xiang et al., 2007). This strain is highly sensitive to all alkali-metal-cations salts and the expression of *Hs*NHA2 improved the growth of these cells in the presence of sodium or lithium in a pH dependent manner (Fuster et al., 2008; Xiang et al., 2007). Examination of the growth of yeast cells expressing mutated variants of *Hs*NHA2 in the presence of salts was also used to identify some residues that are important for the activity of *Hs*NHA2 (Matsuoka et al., 2022; Schushan et al., 2010). However, the transport activity of the native (or mutated) *Hs*NHA2 antiporter(s) has never been measured directly in yeast cells.

In this work, we first optimized the functional expression of *Hs*NHA2 in *S. cerevisiae* cells and obtained a highly valuable experimental model that we next used for *in vivo* determination of effects of *Hs*NHA2 mutations that alter its substrate specificity, transport activity, stability, or are among the single-nucleotide polymorphisms (SNP) in humans. In addition, for the first time, we demonstrate that the hydrophilic N-terminus (and negatively charged residues in this part of the protein) has a regulatory (inhibition) effect on *Hs*NHA2 activity. Altogether, our data suggest an allosteric link between the hydrophilic N-terminus and transmembrane core of the protein.

## Results

### Optimization of *Hs*NHA2 expression in yeast cells

A eukaryotic cell model organism, the yeast *S. cerevisiae*, has been used for functional characterization of human proteins for a long time. Several mammalian Na^+^/H^+^ antiporters were also studied via their expression in *S. cerevisiae* (Flegelova et al., 2006; Flegelova and Sychrova, 2005), including *H. sapiens* NHA2 (Fuster et al., 2008; Matsuoka et al., 2022; Schushan et al., 2010; Xiang et al., 2007). In these studies, *Hs*NHA2 was expressed under the control of a strong promoter (*PGK1* or *TEF1*) from a multi-copy plasmid in a salt-sensitive strain (AB11c; *nha1Δ ena1-4Δ nhx1Δ*). *Hs*NHA2 activity was then observed as an pH-dependent increase in tolerance to sodium or lithium estimated by the growth of cells either in liquid media or on plates containing the corresponding salts (Fuster et al., 2008; Matsuoka et al., 2022; Schushan et al., 2010; Xiang et al., 2007). However, in our first experiment, we realized that using one of these previously made constructs (pTEF1-HsNHA2-HA, in which *Hs*NHA2 is expressed under the control of the *TEF1* promoter (Table1; (Fuster et al., 2008)) is partially toxic for AB11c cells as well as for another strain, BW31, which only lacks the two plasma-membrane Na^+^ exporters (*nha1Δ ena1-4Δ*). In both cases, cells expressing *Hs*NHA2 grew under non-stress conditions significantly slower than cells containing the empty vector (shown for the BW31 strain in Fig. 1, control plates). Therefore, we next carefully examined and optimized the conditions of the expression of *Hs*NHA2 in *S. cerevisiae* to obtain a valuable experimental model useful for the *in vivo* determination of *Hs*NHA2 transport activity. To select an optimal level of expression (non-toxic for yeast cells), we constructed seven new plasmids (centromeric or multi-copy; Table 1), in which the *Hs*NHA2 was expressed under the control of a weak and constitutive *S. cerevisiae NHA1* promoter (Banuelos et al., 2002). Four plasmids contained a GFP sequence added in frame either at the 5’- or 3’-ends of *Hs*NHA2 to confirm a plasma-membrane localization of *Hs*NHA2 in yeast cells (Table 1; (Schushan et al., 2010; Xiang et al., 2007)). Since the presence of a terminator behind any gene is important for mRNA export from the nucleus to the cytoplasm, as well as its stability and translation efficiency (Zhao et al., 1999), an efficient *S. cerevisiae* terminator *TPS1* (Yamanishi et al., 2011) was added behind the *Hs*NHA2 cDNA in four versions of plasmids (Table 1). BW31 cells were transformed with all new plasmids, and we compared the level of cell salt tolerance provided by *Hs*NHA2 expressed from particular plasmids (Fig. 1B). Salt tolerance was tested on YNB-Pro plates with the pH adjusted to 4.0 to ensure the necessary gradient of H^+^ for the transport activity of *Hs*NHA2 (Fuster et al., 2008). Adjustment of the media to lower pH from approximately 4.8 to 4.0 with HCl accentuated the phenotypes described below (results obtained on non-adjusted YNB-Pro plates are not shown). Cells expressing empty vectors (YEp352, pRS316) or *S. cerevisiae* Nha1 antiporter (non-tagged or tagged with GFP) were used as negative or positive controls, respectively. BW31 cells containing an empty vector (YEp352 or pRS316; Table 1) were highly sensitive to all three salts tested (LiCl, NaCl, KCl), on the other hand, cells expressing the *Sc*Nha1 antiporter were able to grow in the presence of LiCl, NaCl, and KCl (Fig. 1B), as *Sc*Nha1 is able to export all three cations from cells (Kinclova et al., 2001). None of the constructs with *Hs*NHA2 improved the cell growth on KCl plates, which confirmed the previous results that *Hs*NHA2 is not transporting K^+^ (Fuster et al., 2008; Kondapalli et al., 2012).

**Table 1:** Plasmids used in this study.

| Plasmid | Description* | Source/Reference |
| --- | --- | --- |
| YEp352 | Multi-copy empty vector | (Hill et al., 1986) |
| pHsNHA2 | *H. sapiens* NHA2 in YEp352 | This work |
| pGRU1 | Multi-copy empty vector with GFP | B. Daignan-Fornier,<br>NCBI Acc. No. AJ249649 |
| pHsNHA2-GFP | *H. sapiens* NHA2 in pGRU1 | This work |
| pGFP-*HsNHA2* | N-terminally GFP tagged *HsNHA2* in YEp352 | This work |
| pHsNHA2t | *H.sapiens* NHA2 with *TPS1* terminator in YEp352 | This work |
| pGFP-*HsNHA2t* | N-terminally GFP tagged *HsNHA2* with *TPS1*<br>terminator in YEp352 | This work |
| pNHA1-985 | *S. cerevisiae* NHA1-985 in YEp352 | (Kinclova et al., 2001) |
| pNHA1-985GFP | *S. cerevisiae* NHA1-985 in pGRU1 | (Kinclova et al., 2001) |
| pTEF1-*HsNHA2*-HA | *H. sapiens* NHA2 in p426TEF | (Fuster et al., 2008) |
| pRS316 | Centromeric empty vector | (Sikorski and Hieter,<br>1989) |
| pC-*HsNHA2t* | *H. sapiens* NHA2 with *TPS1* terminator in pRS316 | This work |
| pC-GFP-*HsNHA2t* | N-terminally GFP tagged *HsNHA2* with *TPS1*<br>terminator in pRS316 | This work |
| YEp352-ZrSTL1-TPS1t-kanMX | plasmid for the expression of *Zygosaccharomyces*<br>*rouxii* *STL1* gene downstream with the *TPS1*<br>terminator | (Duskova et al., 2015) |

\* *ScNHA1* or *HsNHA2* were expressed under the control of *S. cerevisiae* NHA1 promoter, except in pTEF1-*HsNHA2*-HA, in which TEF1 promoter was used.

The expression of *Hs*NHA2 from new plasmids was not toxic for cells, as all transformants grew as well as cells containing empty vectors or expressing the yeast Nha1 antiporter on control plates (and better than cells transformed with pTEF1-HsNHA2-HA) (Fig. 1B). The level of *Hs*NHA2 expression from centromeric plasmids (pC-*Hs*NHA2*t*, pC-GFP-*Hs*NHA2*t*) was too low to observe its activity, since cells with these plasmids grew similarly to cells with empty vectors (Fig. 1B). On the other hand, the expression of *Hs*NHA2 under the control of the *ScNHA1* promoter from new multi-copy plasmids increased the cell tolerance to LiCl and NaCl, although they differed in the level of tolerance provided (Fig. 1B). Cells expressing *Hs*NHA2 from p*Hs*NHA2*t* or p*Hs*NHA2-GFP*t* plasmids were more tolerant to salts than cells with the same constructs without the terminator (Fig. 1B). Thus, it is evident that the use of multi-copy plasmids with a weak *ScNHA1* promoter and *TPS1* terminator was the best for studying the transport properties of human NHA2 expressed in yeast.

To consider various types of expression for the correct targeting of *Hs*NHA2 to the plasma membrane, we first compared the localization of GFP-tagged versions of *Hs*NHA2 in AB11c and BW31 cells, and realized that the lack of intracellular Nhx1 antiporter in the AB11c strain resulted in a higher intracellular stacking of GFP-*Hs*NHA2 than in BW31 cells (Fig. S1). Thus, we subsequently only used the BW31 strain. As shown in Fig. 1C, the GFP-tagging of *Hs*NHA2 at the C-terminus also resulted in a partial stacking of the antiporter in the endoplasmic reticulum (Fig. 1C), and correspondingly, the tolerance of cells expressing *Hs*NHA2-GFP was lower than that of cells with non-tagged *Hs*NHA2 or *Hs*NHA2 tagged with GFP at the N-terminus (Fig. 1B). *Hs*NHA2 with a GFP tag at the N-terminus exhibited similar plasma-membrane localization to *Sc*Nha1-GFP (Fig. 1C). The expression of GFP-*Hs*NHA2 from a plasmid with a *TPS1* terminator (pGFP-*Hs*NHA2*t*) resulted in a diminished level of intracellular fluorescent spots observed in cells containing pGFP-*Hs*NHA2 (Fig. 1C, compare the upper two images).

Thus, from all these tested conditions, for the optimal functional expression of *Hs*NHA2 in yeast cells and further experiments, we selected the BW31 strain transformed either with the p*Hs*NHA2*t* or pGFP-*Hs*NHA2*t* plasmids.

### Point mutations of Pro246 modify the ion selectivity of the antiporter for Na^+^ and/or Li^+^ and its transport activity at pH 7.0

The membrane parts of Na^+^/H^+^ antiporters are highly conserved and are sufficient for ion exchange (Hendus-Altenburger et al., 2014; Kinclova et al., 2001). All CPAs share a similar transmembrane topology, known as the NhaA fold (Hunte et al., 2005) organized into two functional domains – a dimerization domain and a conserved core domain encapsulating the ion-binding site (Matsuoka et al., 2022; Padan, 2014). In the last phylogenetic analysis, a minimal set of eight highly conserved positions in the protein core that are present in all CPAs was identified (Masrati et al., 2018). In *Hs*NHA2, this motif contains, among others, proline 246 in TMS 6 and arginine 432 in TMS 12 (Fig. 1A, Fig. S1). The replacement of an equivalent proline with a polar amino-acid residue (serine or threonine) in the homologous Na^+^, Li^+^/H^+^ antiporter Sod2-22 from *Zygosaccharomyces rouxii* resulted in an ability to recognize and transport K^+^ cations (Kinclova-Zimmermannova et al., 2005). Thus, in the next part of this work, we tested how a substitution of Pro246 in *Hs*NHA2 can change its substrate specificity and transport activity (Fig. 2). Proline 246 was replaced by site-directed mutagenesis either with glycine, a small hydrophobic amino acid (alanine) or polar residues (serine or threonine), and mutated versions were expressed in the BW31 strain from the corresponding p*Hs*NHA2*t* or pGFP-*Hs*NHA2*t* plasmids. Transformants were tested for their tolerance to LiCl, NaCl or KCl at pH 4.0 or 7.0 together with cells transformed with the empty vector or expressing the native *Hs*NHA2 antiporter (Fig. 2A). None of the mutated versions improved the tolerance of cells to KCl (not shown), but we observed that their ability to provide cells with tolerance to LiCl or NaCl within the pH range 4.0 – 7.4 was different from the native antiporter (Fig. 2A, E). The GFP-tagging had no effect on the substrate specificity of mutated versions (cf. below), but it slightly enhanced the phenotypes observed as cell salt tolerance on plates (compare Fig. 2A and Fig. S3A). All four mutated versions were localized to the plasma membrane (Fig. 2C), like the native GFP-*Hs*NHA2 (Fig. 1C), and similar amounts of proteins in cells were detected by immunoblot for all of them (Fig. 2D).

**Figure 2:**
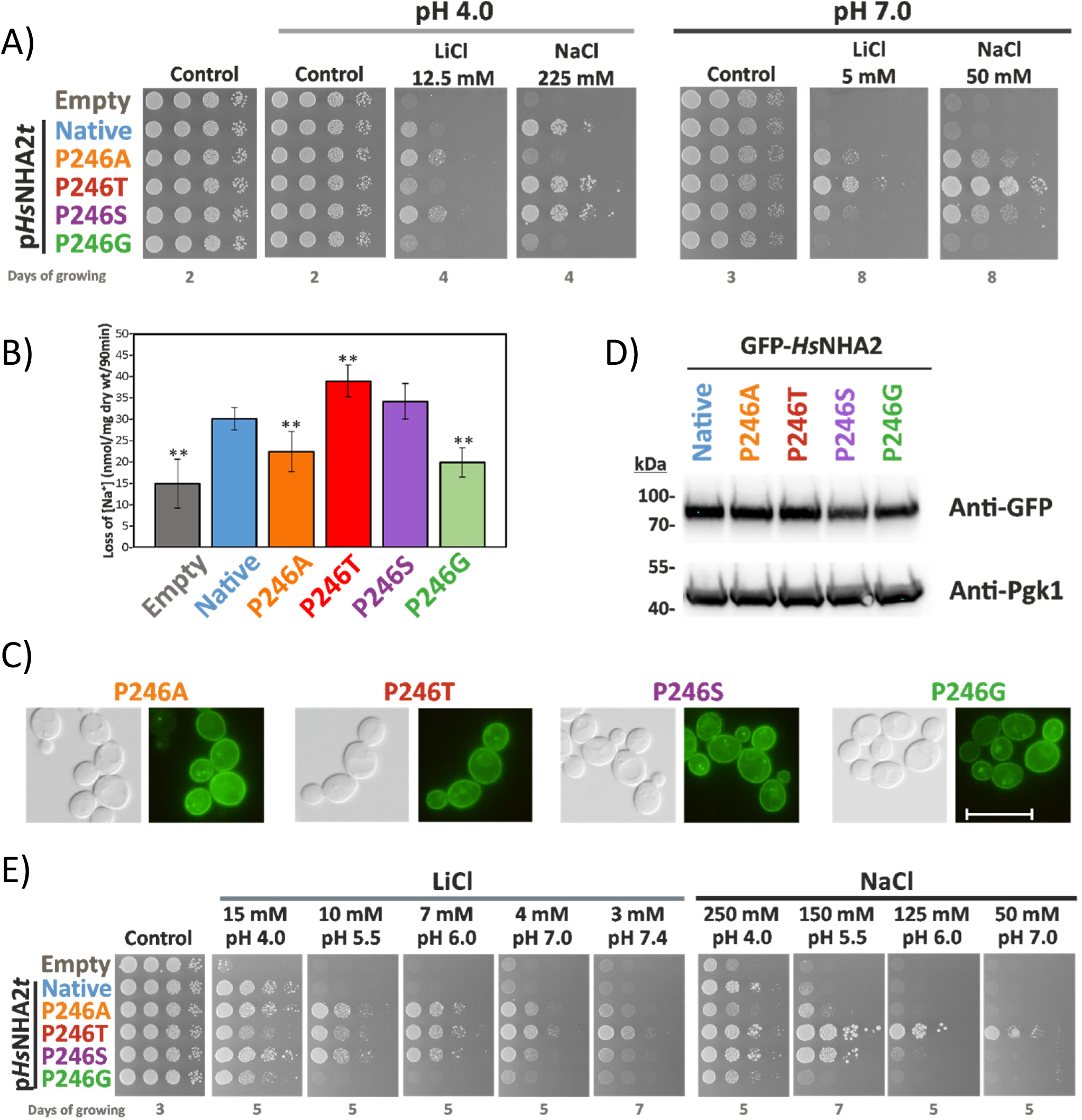
Characterization of *Hs*NHA2 with point mutations at proline 246. A) Salt tolerance of *S. cerevisiae* BW31 cells containing empty vector or expressing native *Hs*NHA2 or one of four *Hs*NHA2 versions with single point mutations P246A, T, S or G from p*Hs*NHA2*t* plasmid. Cells were grown on YNB-Pro or non-buffered YNB-Pro plates with pH adjusted to 4.0 or 7.0 and supplemented with LiCl or NaCl as indicated. Plates were incubated at 30 °C and photographed on the indicated day. B) Loss of Na^+^ from same cells as described in A). Cells were grown in YNB-Pro media, preloaded with Na^+^, transferred to Na^+^-free incubation buffer at pH 4.0 and changes in the intracellular Na^+^ content were estimated as described in the Materials and Methods section. Columns show the amount of Na^+^ lost from cells within 90 min. Data represent the mean of at least 5 replicate values ± SD. Significant differences compared to cells expressing the native *Hs*NHA2 are indicated with asterisks (** *p* < 0.01). Localization (C) and immunodetection (D) of N-terminal GFP-tagged *Hs*NHA2 with single point mutations at Pro246. BW31 cells expressing variants of GFP-*Hs*NHA2 from pGFP-*Hs*NHA2*t* were grown in YNB-Pro (4% glucose) to the exponential phase and observed under a fluorescence microscope (C, right). A Nomarski prism was used for whole-cell imaging (C, left). In D), protein extracts of cells were prepared as described in the Materials and Methods section, subjected to SDS-PAGE (10% gel) and transferred to a nitrocellulose membrane. Native GFP-*Hs*NHA2 and its variants were detected with an anti-GFP antibody. The membrane was reprobed and incubated with an anti-Pgk1 antibody to verify the amount of loaded proteins. E) Activity of mutated *Hs*NHA2 antiporters at various extracellular pH was determined by growth of cells on YNB-Pro plates buffered to various pH levels (4.0 – 7.4) and supplemented with LiCl or NaCl as indicated. Images were taken on the indicated day of incubation at 30 °C.

As expected, the presence of the native antiporter only increased cell tolerance to sodium and lithium at pH 4.0 (when there is a high H^+^ gradient across the membrane), but not at pH 7.0 (Fig. 2A). The P246G substitution resulted in an almost inactive antiporter, as cells with this mutated version grew only slightly better in the presence of salts than cells containing the empty vector (Fig. 2A). Alanine instead of Pro246 resulted in an antiporter which was able to transport Li^+^ better than the native antiporter, but its ability to improve the NaCl tolerance was much lower than that of the native *Hs*NHA2 (Fig. 2A). In contrast, Pro246 replaced with threonine resulted in an antiporter with the opposite substrate preferences to the P246A version, i.e. transporting lithium worse and sodium better than the native *Hs*NHA2 (Fig. 2A). Finally, *Hs*NHA2 with the P246S mutation provided a better tolerance of cells to both cations compared to the native *Hs*NHA2 (Fig. 2A).

Interestingly, and in contrast to the native antiporter, *Hs*NHA2 harbouring the point mutations P246A/S/T also exhibited activity at a higher pH than 4.0. The presence of these mutated versions improved the tolerance of BW31 cells to both cations on non-buffered plates with the pH adjusted to 7.0 (Fig. 2A), where the P246T version provided the most robust growth (Fig. 2A). In the drop test shown in Fig. 2E, we verified this observation on plates with buffered media (c.f. Materials and Methods). On these plates, extracellular pH should not change with the growth of cells, as in Fig. 2A. It is evident that under these conditions, each of the three P246A/T/S versions was able to improve the tolerance of cells to LiCl within the pH range 4.0 – 7.4, and the P246T/S versions also to NaCl (Fig. 2E). Note that in contrast to P246A/S, the ability of the P246T version to improve the tolerance to LiCl increased with increasing extracellular pH (Fig. 2E). This drop test also confirmed the decreased activity of the P246G version in comparison with the native *Hs*NHA2, in which Li^+^ and Na^+^ transport activity was only observable on plates with low pH 4.0 (Fig. 2E or 2A).

Next, the differences in capacity to transport Na^+^ cations observed in drop tests (Fig. 2A, E) were verified by sodium loss measurements at pH 4.0 (Fig. 2B) from sodium-preloaded cells (cf. Materials and Methods). During the experiment, the intracellular concentration of Na^+^ also slightly decreased in control cells without any antiporter (Fig. 2B, empty vector), but significantly higher efflux was observed in cells with the native *Hs*NHA2 (Fig. 2B). In accordance with the drop tests, the amounts of sodium lost from cells expressing the *Hs*NHA2 antiporter with the P246A or P246G mutation were lower than from cells with the native *Hs*NHA2, but higher than from cells with the empty vector (Fig. 2B). On the other hand, the P246S, and especially the P246T mutation increased *Hs*NHA2 transport activity for Na^+^ as the amount of sodium lost from these cells was significantly higher than from cells with the native antiporter (Fig. 2B). Since the localization and amounts of proteins did not change with the P246 substitutions (Fig. 2C, D), sodium loss measurements fully reflected the sodium transport capacity of the particular mutated versions observed as differences in growth on plates with NaCl (Fig. 2A, E).

All these data identified Pro246 as a crucial residue involved in the ability to recognize and transport particular substrates (ion selectivity) and most likely also influencing the capacity to transport protons, as the pH profile of *Hs*NHA2 transport activity changed, especially when the polar-branched amino acid threonine was at this position.

### *Hs*NHA2 with substitutions at Pro246 are less sensitive to phloretin inhibition

The activity of *Hs*NHA2 is insensitive to amiloride (a hallmark of plasma-membrane NHE antiporters), and sensitive to phloretin (Kondapalli et al., 2012), even when it is expressed in yeast cells (Xiang et al., 2007). Phloretin (a flavonoid abundant in apples) is a highly flexible molecule with the ability to bind to biological macromolecules. It has interesting pharmacological and pharmaceutical potential due to its antimicrobial, antioxidant, anti-inflammatory and anticancer activities (Behzad et al., 2017). The mechanism of *Hs*NHA2 inhibition by phloretin is unclear. To determine whether mutations of Pro246 change the sensitivity of the antiporter to phloretin, we tested the growth of cells expressing the native or either of the four mutated antiporters with a substitution at proline P246 on plates supplemented with 15 mM LiCl or 250 mM NaCl and increasing concentrations of phloretin (Fig. 3).

**Figure 3:**
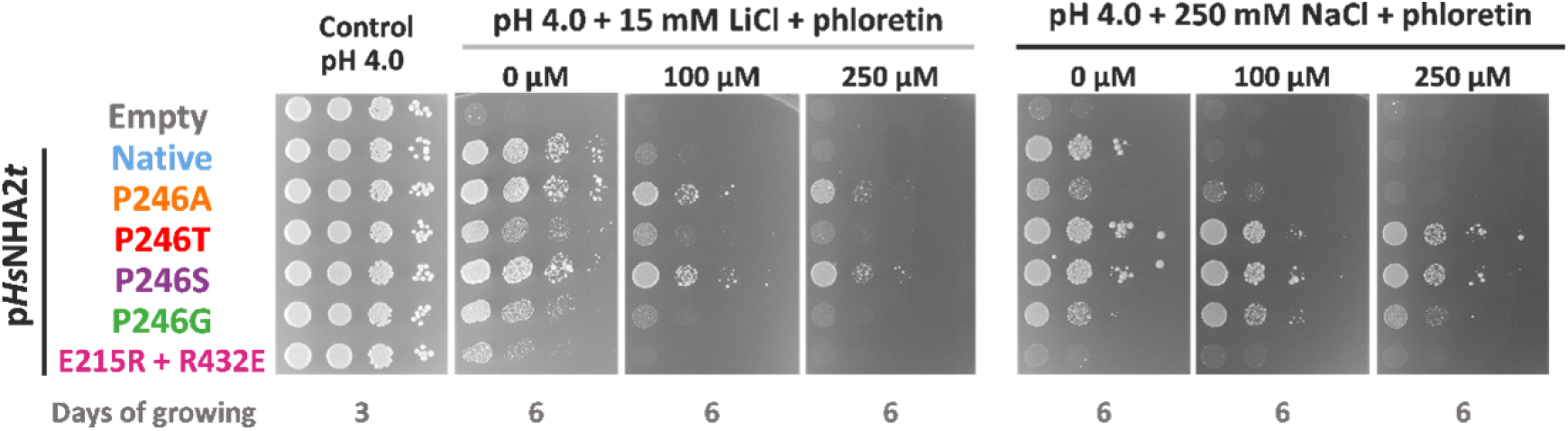
*Hs*NHA2 variants of proline 246 are less sensitive to phloretin inhibition. *S. cerevisiae* BW31 cells containing empty vector or expressing native *Hs*NHA2 or *Hs*NHA2 versions with point mutations P246A, T, S or G or the double mutation E215R + R432E from p*Hs*NHA2*t* plasmid, as indicated, were used in this experiment. The inhibition of mutated *Hs*NHA2 versions by phloretin was determined by the growth of cells on non-buffered YNB-Pro plates with the pH adjusted to 4.0 and supplemented with LiCl or NaCl, and with or without phloretin at the indicated concentrations. Plates were incubated at 30 °C for 3 (controls) or 6 days (phloretin).

In accordance with previous results (Xiang et al., 2007), the ability of the native *Hs*NHA2 to improve LiCl tolerance at pH 4.0 of yeast cells highly decreased in the presence of phloretin (Fig. 3). Here we also demonstrate the same effect of phloretin on Na^+^ transport activity (Fig. 3). Remarkably, the inhibition effect of phloretin decreased when proline 246 was replaced with alanine/glycine or even disappeared when serine or threonine was at this position (Fig. 3). BW31 cells transformed with these versions of *Hs*NHA2 grew on plates with LiCl or NaCl and phloretin very similarly to when the inhibitor was not present (Fig. 3).

These results reinforced the importance of proline 246 for the proper activity of *Hs*NHA2. They also indicate that this residue could take part in a mechanism of inhibition mediated by phloretin.

### Swapping E215 and R432 results in a Li^+^-specific antiporter that is also active at higher pH

As was mentioned above, R432 is one of the eight highly conserved amino acids of the whole CPA family, and it was predicted to potentially form a salt bridge with E215 in TMS 5 (Fig. 1A; (Masrati et al., 2018). The recently released cryo-EM structures of N-terminally truncated human (PDB 7B4M/L) or bison NHA2 confirmed the proximity of these two residues (Matsuoka et al., 2022). To experimentally prove the importance and function of these two titratable residues, we prepared three mutated versions of *Hs*NHA2 in which we either introduced the opposite charge at these positions (E215R or R432E versions) or we swapped these two residues (E215R + R432E double mutant). The effect of mutation(s) on the transport properties of *Hs*NHA2 was tested similarly as above.

The drop test in Fig. 4A shows that the single-mutated versions of the antiporter, E215R or R432E, were non-functional. Cells transformed with p*Hs*NHA2*t* (E215R) or p*Hs*NHA2*t* (R432E) plasmids did not grow in the presence of NaCl or LiCl (at both pHs, 4.0 or 7.0), just like cells containing the empty vector (Fig. 4A). On the other hand, swapping these two residues led to a functional antiporter, however with substrate specificity limited to Li^+^ cations. At both of the tested pH levels, cells expressing *Hs*NHA2 (E215R + R432E) were only able to grow on plates with LiCl, but not with NaCl (Fig. 4A). Similar results were observed in a drop test with cells expressing the same mutated versions, but tagged with GFP at the N-terminus (Fig. S3B).

**Figure 4:**
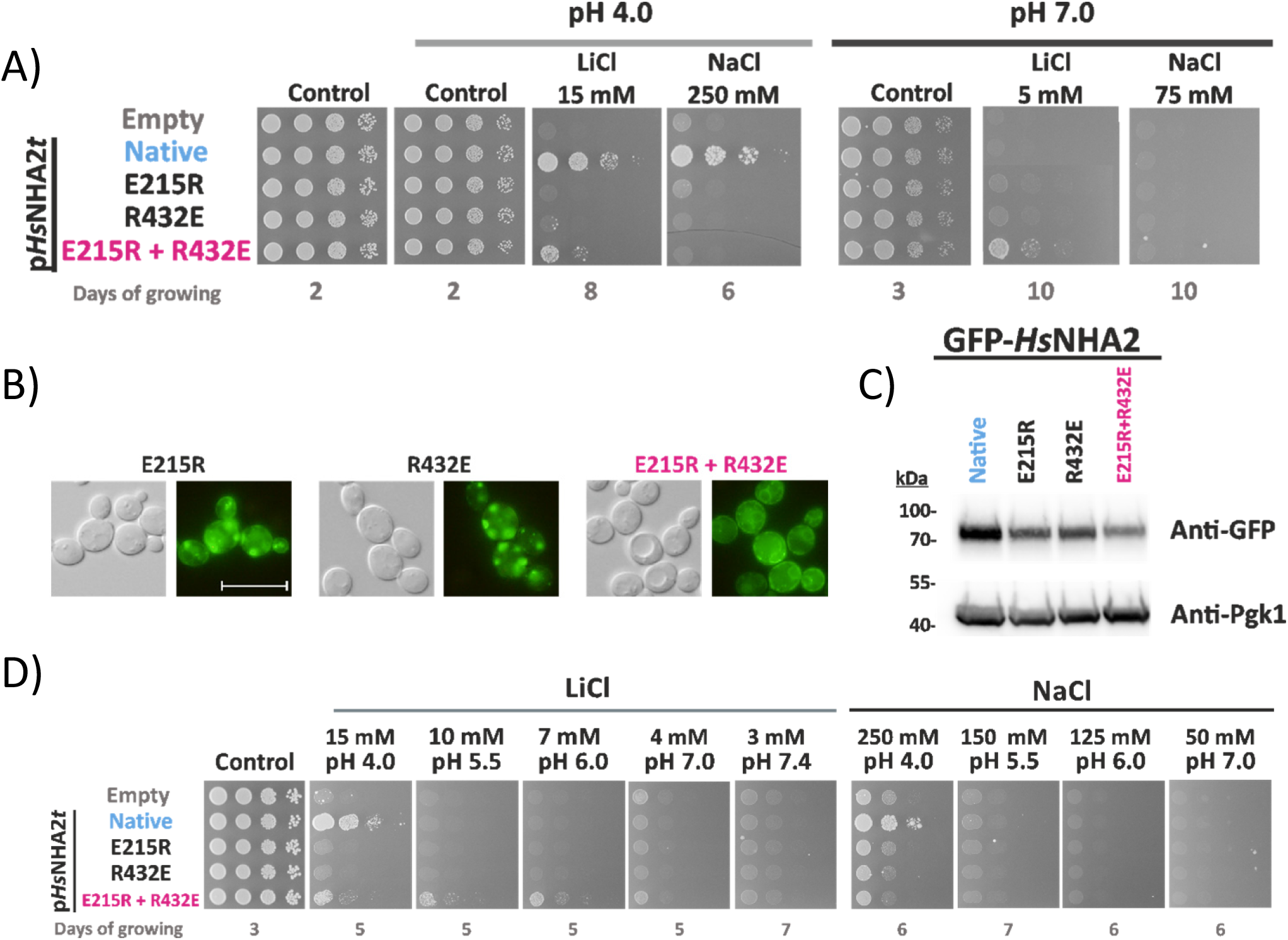
Characterization of *Hs*NHA2 with mutations of residues forming a putative salt bridge. A) Salt tolerance of *S. cerevisiae* BW31 cells containing empty vector or expressing native *Hs*NHA2 or its mutated versions (E215R, R432E or E215R + R432E) from p*Hs*NHA2*t* plasmid. Cells were grown on YNB-Pro (pH approx. 4.8) or non-buffered YNB-Pro plates with the pH adjusted to 4.0 or 7.0 and supplemented with LiCl or NaCl as indicated. Plates were incubated at 30 °C and photographed on the indicated day. Localization (B) and immunodetection (C) of N-terminal GFP-tagged *Hs*NHA2 mutated versions E215R, R432E or E215R + R432E. Cells were grown in YNB-Pro (4% glucose) to the exponential phase and observed under a fluorescence microscope (B, right). A Nomarski prism was used for whole-cell imaging (B, left). The scale bar corresponds to 10 μm. In C), protein extracts from the same cells as in B) (with cells expressing native GFP-*Hs*NHA2 as a control) were prepared as described in the Materials and Methods section, subjected to SDS-PAGE (10% gel) and transferred to a nitrocellulose membrane. GFP-*Hs*NHA2 was detected with an anti-GFP antibody. The membrane was reprobed and incubated with an anti-Pgk1 antibody to verify the amount of loaded proteins. D) Activity of mutated *Hs*NHA2 antiporters at various extracellular pH levels determined by growth of cells on YNB-Pro plates buffered to various pH levels (4.0 – 7.4) and supplemented with LiCl or NaCl as indicated. Images were taken on the indicated day of incubation at 30 °C.

The loss of function of the two single-mutated versions most likely resulted from their mislocalization in intracellular compartments, observed as fluorescent spots inside cells (Fig. 4B). In contrast, the double-mutated E215R + R432E version tagged with GFP was partially targeted to the plasma membrane and partially observed in the perinuclear ER (Fig. 4B). The level of proteins expressed in cells checked by immunoblotting revealed that all three mutated variants were present in cells in lower amounts than the native antiporter (Fig. 4C). Since all proteins are expressed under the same conditions (promoter, plasmid), it indicate that these mutations affected the structure, stability and plasma-membrane targeting of *Hs*NHA2.

The drop test on plates with salts and pH buffered to various values (Fig. 4D) confirmed the lithium substrate specificity of the E215R + R432E mutated version. In contrast to the native antiporter, the double-mutated *Hs*NHA2 was able to improve the LiCl tolerance of cells not only at pH 4.0, but also at higher pH, up to 6.0 (Fig. 4D). Nevertheless, in contrast to P246 variants, phloretin inhibited the Li^+^ transport ability of the double mutant E215R + R432E, as observed for the native *Hs*NHA2 (Fig. 3).

All these results confirmed E215R and R432E to be crucial residues for *Hs*NHA2 functionality. The transport activity of the antiporter with the E215R mutation in TMS 5 could be restored by introducing a second mutation R432E in TMS 12. Thus, we experimentally confirmed that in the full version of *Hs*NHA2, these two residues are spatially close to each other with a salt bridge between them. We also discovered that the interaction between these two residues is important for the structural integrity of the protein.

### Validation of the system for studying the effect of SNPs in *Hs*NHA2

As was mentioned above, *Hs*NHA2 is associated with a growing list of various pathologies. For this reason, we think that our optimized expression system of *Hs*NHA2 in yeast cells could be useful in the future for the characterizing the effects of a broad range of mutations that are present in human genomes. To test this possibility, we selected six point mutations (G238R, V240L, D278G, R432Q, K382E and A406E) that belong to known human SNPs from patients (Dewey et al., 2016). The positions of these residues in *Hs*NHA2 are shown in Figs. 1A, S1 and 8B, E. Mutated versions were prepared by site-directed mutagenesis and expressed from p*Hs*NHA2t or pGFP-*Hs*NHA2t plasmids in BW31 cells, and we tested the tolerance of cells to LiCl and NaCl at pH 4.0 (Fig. 5A, S4). The localization of the corresponding GFP-tagged versions was observed by fluorescence microscopy (Fig. 5B).

**Figure 5:**
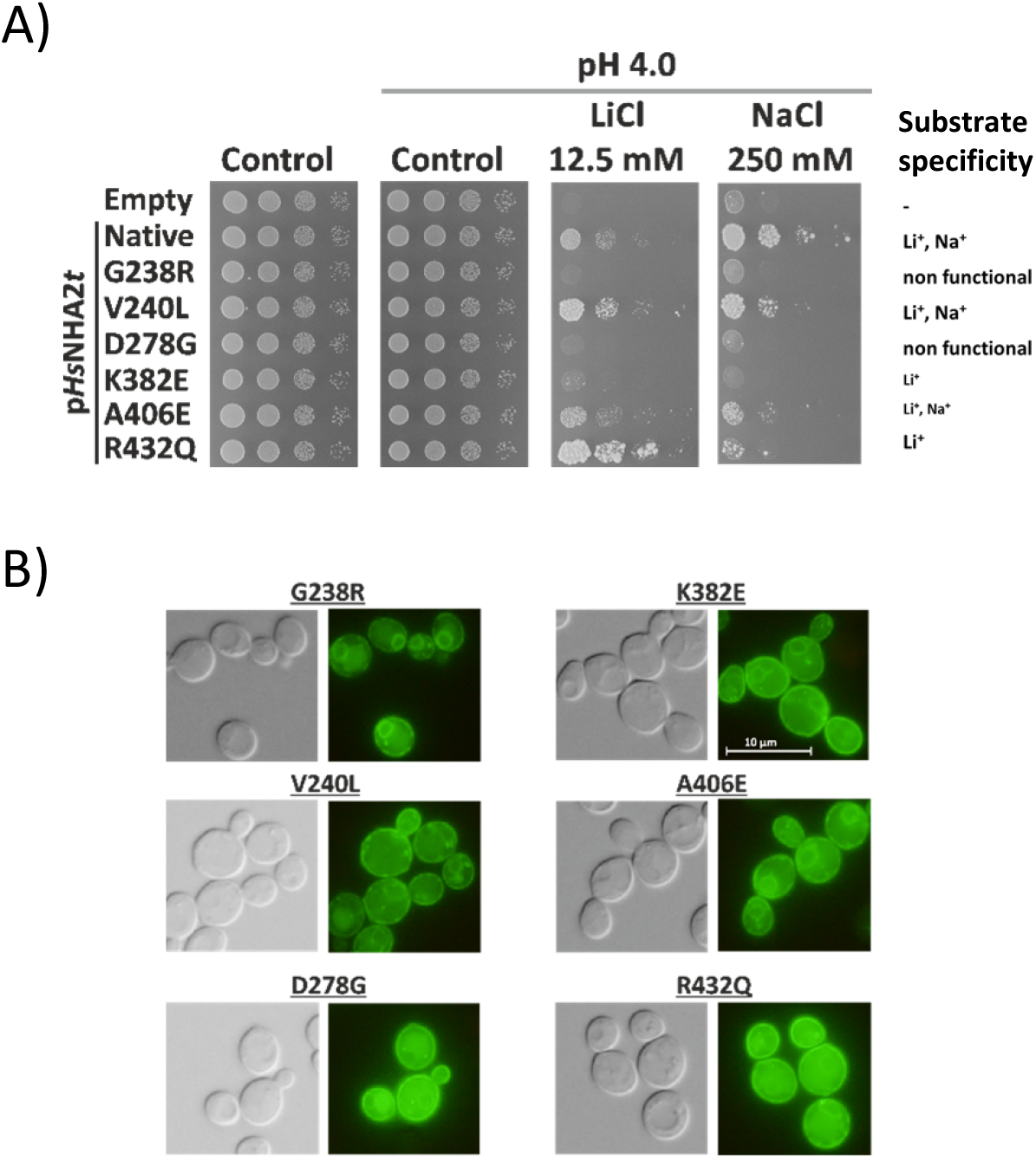
Phenotypes of SNPs of *Hs*NHA2. A) Salt tolerance of *S. cerevisiae* BW31 cells containing empty vector or expressing native *Hs*NHA2 or one of six *Hs*NHA2 mutated versions – G238R, V240L, D278G, K382E, A406E and R432Q from the p*Hs*NHA2*t* plasmid. Cells were grown on YNB-Pro or non-buffered YNB-Pro plates with pH adjusted to 4.0 and supplemented with LiCl or NaCl as indicated. Growth was monitored for 2 (controls) or 7 (LiCl or NaCl) days at 30 °C. B) Localization of N-terminal GFP-tagged *Hs*NHA2 versions. BW31 cells expressing GFP-*Hs*NHA2 with single point mutations as in A) from pGFP-*Hs*NHA2*t* were grown in YNB-Pro (4% glucose) to the exponential phase and observed under fluorescence microscope (B, right). A Nomarski prism was used for whole-cell imaging (B, left). The scale bar corresponds to 10 μm.

All six mutations influenced the transport properties of *Hs*NHA2, but to various extents. The mutation V240L resulted in an antiporter with a preference for transporting Li^+^ over Na^+^, while the R432Q or K382E versions were only able to improve the tolerance to Li^+^, but not to Na^+^, indicating that these mutations narrowed the substrate specificity of the antiporter (Fig. 5A; for K382E cf. also Fig. S4). The point mutation A406E resulted in a less active antiporter for both cations, as cells with this version grew worse in the presence of both salts than cells with the native one (Fig. 5A). Neither of the mutated versions G238R or D278G were functional, because cells with these antiporters grew very similarly in the presence of salts to cells with the empty vector (Fig. 5A). Interestingly, we observed that the non-functionality of G238R resulted from misfolding of the antiporter (stacking in the ER; Fig. 5B), while for D278G, which was localized in the plasma membrane (Fig. 5B), it was instead due to the effect of the mutation on activity (the mechanism of transport). The fluorescence in cells expressing GFP-*Hs*NHA2 with V240L, K382E, A406E or R432Q was also observed predominantly in the plasma membrane (Fig. 5B).

These results confirmed that in the future our expression system may provide an efficient screening of the effects of mutations present in *Hs*NHA2 versions associated with pathologies. In addition, this system can also be useful for the large-scale testing of various compounds that activate/inhibit the *Hs*NHA2 activity.

### Amino acids 1-40 of the hydrophilic N-terminus inhibit the transport activity of *Hs*NHA2

*Hs*NHA2 and its homologs from other mammals possess an unusually long hydrophilic N-terminus (about 80 aa), which is not present in the NHE clade (Masrati et al., 2018; Matsuoka et al., 2022). The structure of the N-terminus is predicted to be disordered (Alpha-fold 2 prediction Q86UD5; (Varadi et al., 2022). In the above drop test experiments, we always noticed that N-terminally GFP-tagged versions of *Hs*NHA2 provided BW31 cells with a higher tolerance to LiCl and NaCl (Fig. S3, S4, S5) than the corresponding variants without the tag (Fig. 2, 4, 5), even though the expression conditions were the same. This observation indicated that GFP-tagging somehow activated *Hs*NHA2, and that the hydrophilic N-terminal part of the protein may play a role in the regulation of the transport activity of *Hs*NHA2 similarly to e. g. hydrophilic C-termini of human NHEs (Hendus-Altenburger et al., 2014) or *S. cerevisiae* Nha1 (Kinclova et al., 2001). In addition, the cryo-EM structure of bison NHA2 truncated by 69 amino acids from the N-terminus indicated the presence of an unusual short helical segment (corresponding to amino acids 69-79) prior to TMS 1 (Matsuoka et al., 2022).

To understand the role of the N-terminus in *Hs*NHA2’s function, we constructed a series of step-by-step truncated *Hs*NHA2 versions, ranging from the complete antiporter down to the shortest version lacking the first 90 amino acids (Fig. S1). In eight newly prepared constructs of *Hs*NHA2, codons for amino acids at positions 20, 40, 60, 70, 75, 80, and 90 were replaced by the start codon ATG for methionine. Truncated *Hs*NHA2 versions were expressed in BW31 cells, and tested for their substrate specificity and transport activity on plates with salts and the pH adjusted to 4.0 or 7.0 (Fig. 6). For each version, a corresponding N-terminal GFP-tagged construct was used to test the localization and level of expression of the encoded protein.

**Figure 6:**
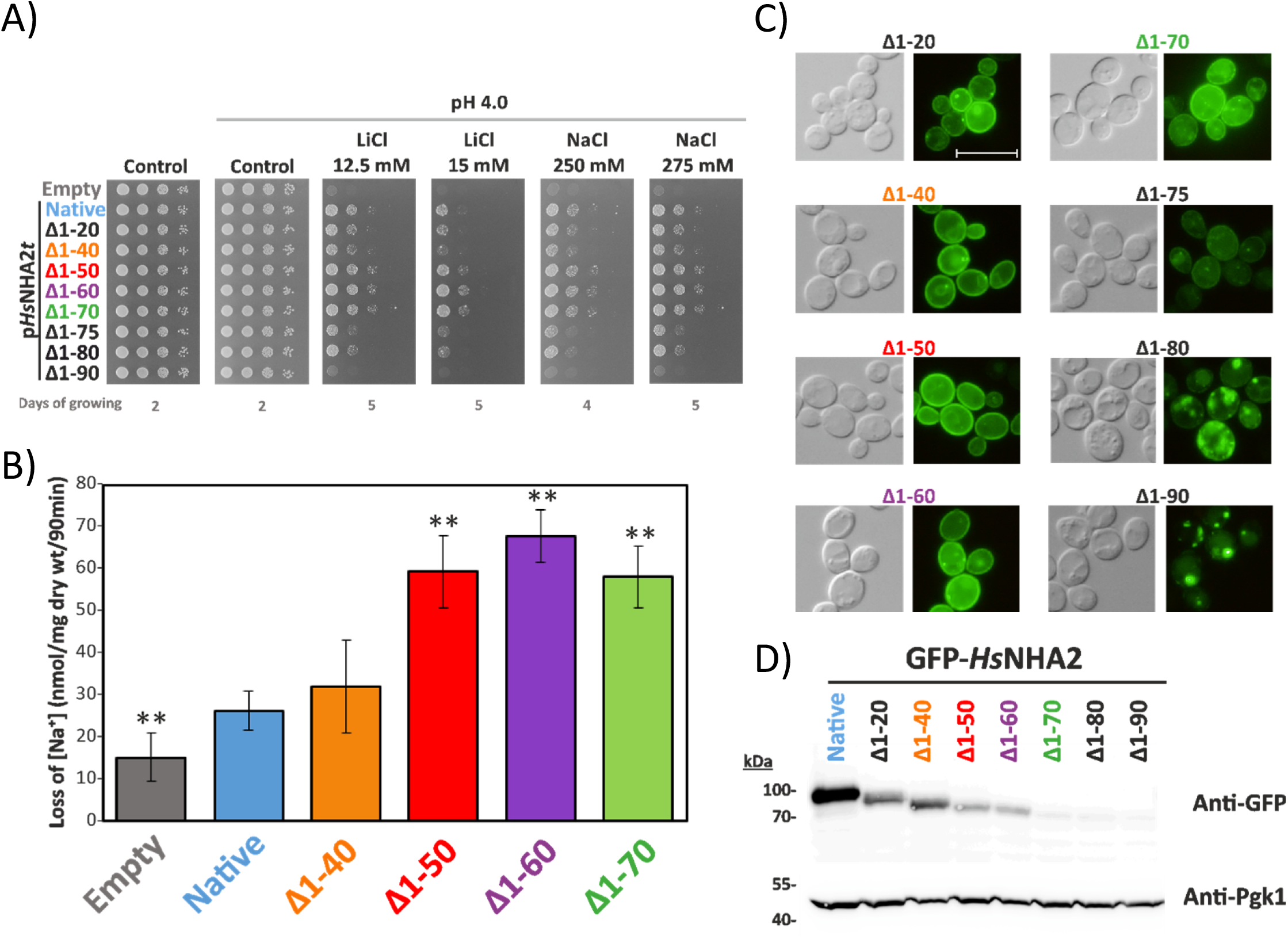
Hydrophilic N-terminus of *Hs*NHA2 plays inhibitory role. A) Salt tolerance of *S. cerevisiae* BW31 cells containing empty vector or expressing native or N-terminally truncated *Hs*NHA2 versions from plasmids derived from p*Hs*NHA2t. Cells were grown on YNB-Pro (pH aprox. 4.8) or non-buffered YNB-Pro plates with pH adjusted to 4.0 and supplemented with LiCl or NaCl as indicated. Plates were incubated at 30 °C and photographed on the indicated day. B) Loss of Na^+^ from BW31 cells mediated by N-terminal truncated *Hs*NHA2 versions. Cells were grown in YNB-Pro media, preloaded with Na^+^, transferred to Na^+^-free incubation buffer at pH 4.0, and changes in the intracellular Na^+^ content were estimated as described in the Materials and Methods section. Columns show the amount of Na^+^ lost from cells within 90 min. Data represent the mean of at least 3 replicate values ± SD. Significant differences compared to cells expressing the native *Hs*NHA2 are indicated with asterisks (** *p* < 0.01). Localization (C) and immunodetection (D) of N-terminal GFP-tagged truncated *Hs*NHA2 versions. BW31 cells expressing variants of GFP-*Hs*NHA2 with truncated N-terminus from plasmids derived from pGFP-*Hs*NHA2*t* were grown in YNB-Pro (4% glucose) to the exponential phase and observed under a fluorescence microscope (C, right). A Nomarski prism was used for whole-cell imaging (C, left). The scale bar corresponds to 10 μm. In D), protein extracts from the same cells as in C) (with cells expressing native GFP-*Hs*NHA2 as a control) were prepared as described in the Materials and Methods section, subjected to SDS-PAGE (10% gel) and transferred to a nitrocellulose membrane. GFP-*Hs*NHA2 variants were detected with an anti-GFP antibody. The membrane was reprobed and incubated with an anti-Pgk1 antibody to verify the amount of loaded proteins.

The lack of the first 40 amino acids had no effect on *Hs*NHA2 transport properties. Cells with Δ1-20 and Δ1-40 truncated versions provided cells with the same salt tolerance as the native *Hs*NHA2 (Fig. 6A). The lack of the first 40 amino acids also did not change the Na^+^-efflux activity of the antiporter (Fig. 6B). On the other hand, cells expressing Δ1-50, Δ1-60 or Δ1-70 versions grew better on LiCl and NaCl plates than cells expressing the native antiporter (Fig. 6A). This suggested that truncation of the first 50 – 70 aa increased the activity of the antiporter, which we confirmed by measurements of Na^+^-efflux activity (Figure 6B). Notably, the amount of Na^+^ lost from cells expressing the Δ1-50, Δ1-60 or Δ1-70 *Hs*NHA2 variants was double that of the native antiporter or Δ1-40 version (Fig. 6B). Truncations longer than 75 amino acids resulted in less efficient transporters, as they provided cells with lower LiCl and NaCl tolerance than the native antiporter, and the removal of 90 amino acids resulted in a non-functional protein (Fig. 6A). Truncations up to 70 amino acids did not influence the plasma-membrane localization of the antiporter (Fig. 6B). In contrast, the fluorescence signal in cells with the Δ1-75, Δ1-80 or Δ1-90 GFP-*Hs*NHA2 antiporters was predominantly observed in intracellular compartments (Fig. 6C). Thus, these versions were probably not folded properly, which led to their mislocalization. Neither of these truncated versions was able to improve the salt tolerance of BW31 cells on plates with pH 7.0 (results not shown), which means that the dependency on the H^+^ gradient of *Hs*NHA2 did not change with N-terminal truncations.

The immunodetection of truncated versions (Fig. 6D) showed that the signal of the GFP-*Hs*NHA2 proteins gradually decreased with larger truncations of the N-terminus, being the weakest for the last three (Δ1-70, Δ1-80 or Δ1-90). This indicates that the N-terminus is not only important for the regulation of transport activity, but also for the stability of the protein and/or its trafficking to the plasma membrane. Interestingly, in contrast to the point mutations studied above (localized within the TMS), the GFP-tagging decreased the ability of truncated *Hs*NHA2 to improve the tolerance of cells to LiCl, but the provided tolerance to NaCl was enhanced with a truncation longer than 40 aa, as with non-tagged antiporters (compare Fig. 6A and S5).

Although the protein signals detected by immunoblotting for the Δ1-50, Δ1-60 or Δ1-70 versions were much lower than that for the native antiporter (Fig. 6D), they exhibited at least double the Na^+^ efflux activity (Fig. 6B). Since we observed this phenomenon in yeast cells, which may lack some regulatory mechanisms or protein interactions that regulate the activity of *Hs*NHA2 in mammalian cells (Anderegg et al., 2021), the first 40-50 amino acids of the protein instead perform an autoinhibitory function in *Hs*NHA2.

### Conserved charged amino acids in the hydrophilic N-terminus of *Hs*NHA2 are important for its activity

It was found that in the hydrophilic N-terminus of the Na^+^/HCO_3_^−^co-transporter, the presence of charged (positively as well as negatively) residues is essential to fulfilling the autoinhibitory function of the transporter (Su et al., 2021). To corroborate the role of the N-terminus in the regulation of *Hs*NHA2 activity, we compared the sequences of unique homologs of NHA2 from various metazoan species (Fig. 7A). In the important region revealed above (amino acids 40 - 70), we identified four charged residues (3 of them highly conserved (Fig. 7A)): negatively charged glutamates 47 and 56, and positively charged lysines at positions 57 and 58), and we estimated their possible role in *Hs*NHA2 activity. We replaced these amino acids with neutral alanine or with an amino acid with the opposite charge. The resulting six mutated versions of *Hs*NHA2 (E47A/K, E56A/K, K57A + K58A, K57E + K58E) were tested for their ability to improve the salt tolerance of BW31 cells at pH 4.0 in comparison to the native antiporter (Fig. 7A). Fluorescence microscopy of GFP-tagged versions revealed that none of these mutations changed the plasma-membrane localization of the antiporter (Fig. 7C), or the amount of proteins in cells (Fig. 7D).

**Figure 7:**
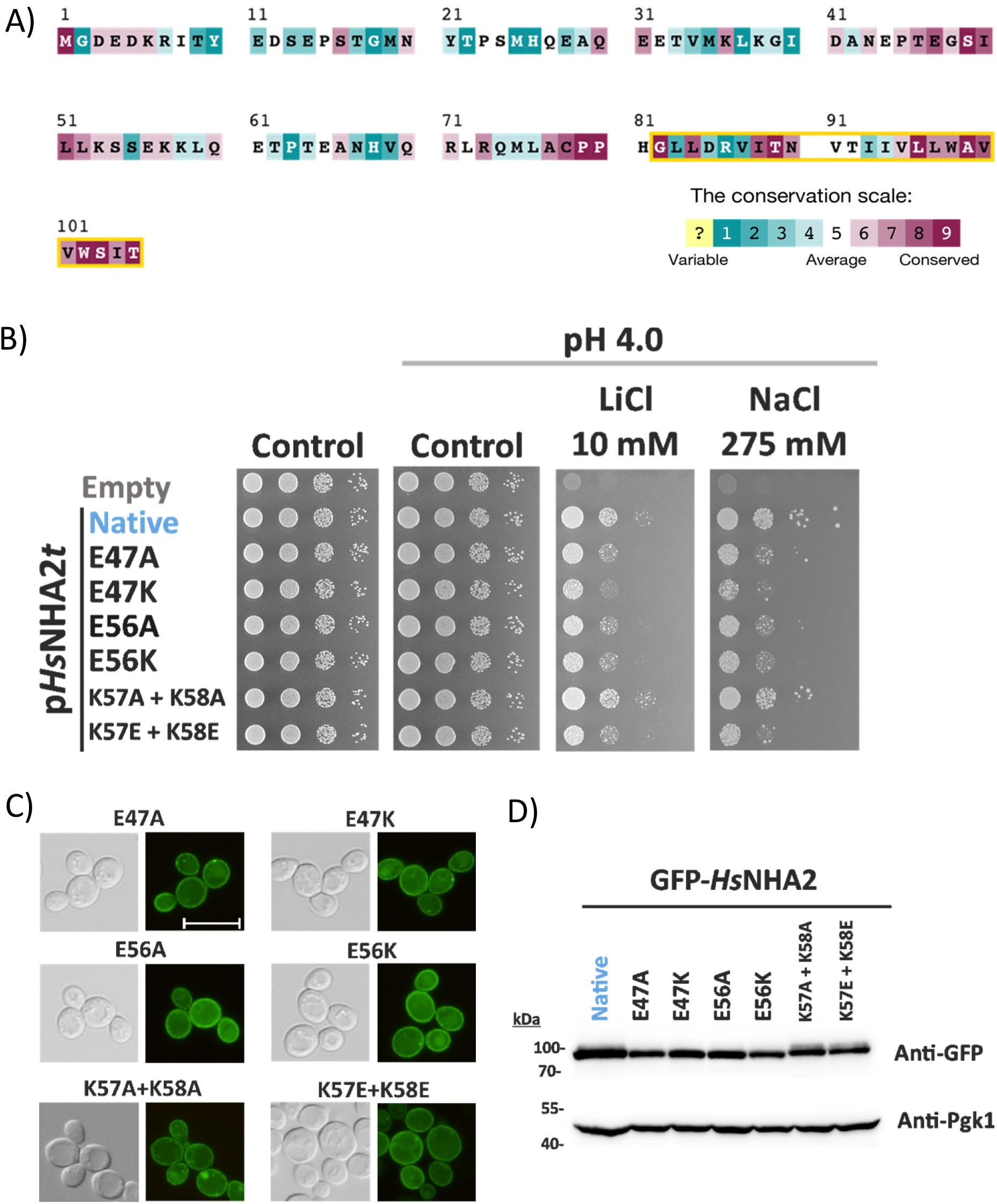
Charged residues in *Hs*NHA2 hydrophilic N-terminus are important for transport activity. A) Evolutionary conservation analysis of *Hs*NHA2 N-terminus. *Hs*NHA2 N-terminal residues 1-to-105 colored based on ConSurf server evolutionary conservation scale, with cyan to maroon representing variable to conserved amino acids, respectively. The position of the unique N-terminal TMS is highlighted with a yellow box. The conservation analysis was conducted using *Hs*NHA2 N-terminal amino acids. A total of 99 unique sequences were collected by HMMER from the UniRef90 database with sequence identity ranging from 35% to 95% and an E-value of at least 0.00001. B) Salt tolerance of *S. cerevisiae* BW31 cells containing empty vector or expressing native *Hs*NHA2 or one of six *Hs*NHA2 mutated versions E47A, E47K, E56A, E56K, K57A + K58A or K57E + K58E from p*Hs*NHA2*t.* Cells were grown on YNB-Pro or non-buffered YNB-Pro plates with the pH adjusted to 4.0 and supplemented with LiCl or NaCl as indicated. Plates were incubated at 30 °C for 2 (controls) or 5 days (salts). Localization (C) and immunodetection (D) of N-terminal GFP-tagged *Hs*NHA2 mutated versions as in B). BW31 cells expressing variants of GFP-*Hs*NHA2 from the pGFP-*Hs*NHA2*t* were grown in YNB-Pro (4% glucose) to the exponential phase and observed under a fluorescence microscope (C, right). A Nomarski prism was used for whole-cell imaging (C, left). The scale bar corresponds to 10 μm. In D), protein extracts from the same cells as in C) were prepared as described in the Materials and Methods section, subjected to SDS-PAGE (10% gel) and transferred to a nitrocellulose membrane. GFP-*Hs*NHA2 variants were detected with an anti-GFP antibody. The membrane was reprobed and incubated with an anti-Pgk1 antibody to verify the amount of loaded proteins.

The drop test (Fig. 7B) showed that both glutamate residues E47 and E56 are important for the antiporter’s activity. Their replacement with alanine resulted in antiporters with a lower ability to improve the lithium and sodium tolerance of cells, and their replacement with an amino-acid with the opposite charge (lysine) emphasized this phenotype (Fig. 7B). The replacement of K57 and K58 with alanines did not affect the antiporter’s function, but negatively charged glutamates at these two positions also resulted in an antiporter with decreased activity (Fig. 7B).

In summary, these experiments revealed a new role of the N-terminus in the regulation of *Hs*NHA2 activity and identified four conserved charged amino-acid residues in the N-terminus as crucial for determining *Hs*NHA2 transport activity.

## Discussion

As a unicellular eukaryotic organism, the yeast *S. cerevisiae* plays a key role in fundamental scientific research in exploring molecular aspects of cellular processes. Thanks to the identification of human orthologs to yeast proteins, including proteins related to human pathologies, yeast is widely used as a simplified cellular model of human diseases (Thines et al., 2019). The heterologous expression of human proteins in yeast cells also provides valuable information about the structure of human proteins and their functional domains (Stribny et al., 2020). We optimized the expression of the human Na^+^/H^+^ antiporter NHA2 in yeast cells to be stable, functional, at a non-toxic level for cells, and with proper targeting to the plasma membrane (Fig. 1). For the first time, we set the conditions and methodology for *in vivo* measurements of its Na^+^ transport activity (Figs. 2 and 6). Using this system enabled us to identify several as-yet unknown residues crucial for *Hs*NHA2 substrate specificity and transport activity (Figs. 2, 4, 5, 7), and to discover a possible autoinhibitory role of the hydrophilic N-terminal segment (Figs. 6 and 7), which appears to allosterically regulate transmembrane ion transport. Our experimental model is also a key tool for better understanding the molecular causes of pathologies related to single-nucleotide polymorphism of *Hs*NHA2, and to screen for putative modifiers, including drugs, which could alter *Hs*NHA2 activity (similarly as is shown in Fig. 5 and 3, respectively).

All Na^+^/H^+^ antiporters seems to share a similar transmembrane topology, known as the NhaA fold, organized into two functional domains - a dimerization domain and a conserved core domain encapsulating the ion-binding site (Hunte et al., 2005; Padan, 2014). According to recently released N-terminally truncated cryoEM structures of two NHA2 proteins (human (PDB 7B4L; Fig. 8A), bison (Matsuoka et al., 2022)), the antiporter consists of 14 TMS, instead of the 13 TMS observed in mammalian NHE Na^+^/H^+^ exchangers (Dong et al., 2021; Winklemann et al., 2020) and the prokaryotic NhaP1, NapA and NhaP antiporters (Goswami et al., 2011; C. Lee et al., 2013; Paulino et al., 2014; Wohlert et al., 2014) or the 12 TMS in *Escherichia coli* antiporter NhaA (Hunte et al., 2005). *Hs*NHA2 possesses a long hydrophilic N-terminus that resides in the cytosol (Fig. 1A, S1), which is followed by an N-terminal transmembrane helix corresponding to amino acids 82 - 105 (Fig. 7A, 8A). Predicted to be unstructured, the hydrophilic N-terminal part preceding TMS1 is naturally missing from both known NHA2 structures that were recently released (Protein Data Bank 7B4M/L; (Matsuoka et al., 2022)). Specifically, in structural studies of bison NHA2, the protein was shortened of 69 amino acids at the N-terminus to reduce predicted disorder (Matsuoka et al., 2022). We have found that this region is important for the regulation of *Hs*NHA2 activity (Fig. 6). Our data show that, in *Hs*NHA2, molecular dissection of this region (aa 1-70) neither affected protein localization (Fig. 6C) nor substrate specificity (Fig. 6A). However, truncation of the first 50 - 70 aa from the N-terminus provided cells with higher tolerance to LiCl and NaCl (Fig. 6A) and resulted in increased Na^+^-transport activity (Fig. 6B). The activity of various Na^+^/H^+^ antiporters, members of the CPA1 clade, was found to be mainly regulated via their large C-terminal hydrophilic domains. A number of phosphorylation sites and binding sites that interact with regulatory proteins were found in these C-terminal segments (Hendus-Altenburger et al., 2014; Kinclova et al., 2001; Pedersen and Counillon, 2019; Smidova et al., 2019). Nevertheless, our results revealed a novel and unique autoinhibitory function of the N-terminus among eukaryotic Na^+^/H^+^ antiporters. So far, an autoinhibitory role of the hydrophilic N-terminus has been described in some other transporters, e.g. the human Ca^2+^ and Mn^2+^ transporter TMEM165 (Stribny et al., 2020) or Na+/HCO3-co-transporter NBCe1-B (Su et al., 2021), plant Ca2+/H+ antiporters (Pittman et al., 2002) or yeast Na^+^, K^+^/H^+^ antiporter Vnx1 (Cagnac et al., 2020).

**Figure 8:**
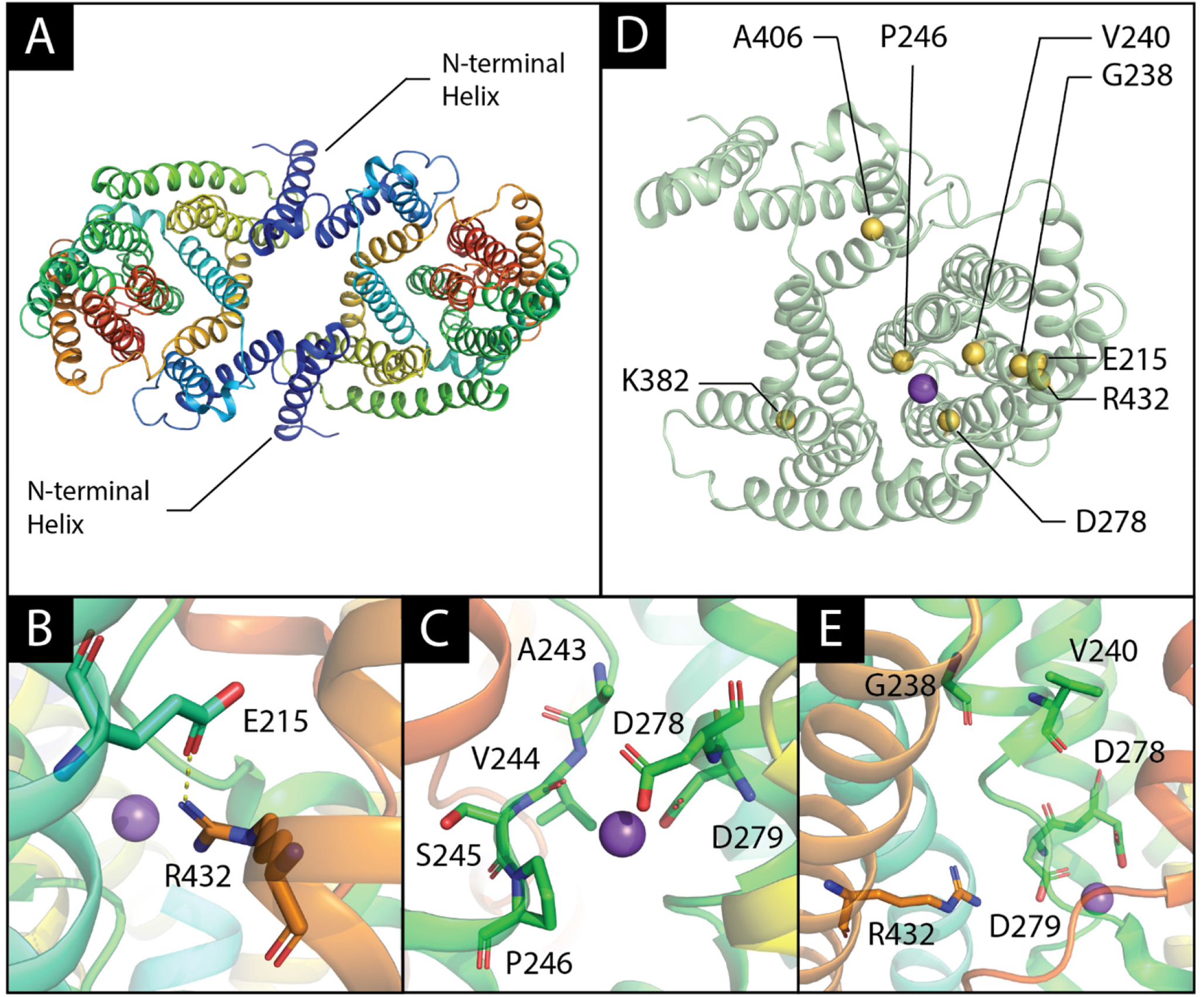
*Hs*NHA2 3D-structure and location of various point mutations. (A) Structure of *Hs*NHA2 dimer, residues 79 - 521 (PDB entry 7B4L) in cartoon representation with rainbow coloring corresponding to N-through C-terminus. The additional N-terminal helix that is missing in other mammalian CPA members with known structures (dark blue) is highlighted. (B, C, E) Close-up view of studied *Hs*NHA2’s point mutations in vicinity of ion-binding site. (B) Previously predicted E215-R432 salt bridge was recently confirmed by newly published structures of both human and bison NHA2, and is likely to play a structural role. (C) P246 is located at unwound section of TMS 6. Its surroundings are shown as sticks, and the bound sodium ion is shown as a purple sphere. Notably, the addition of a polar side-chain in the vicinity of D278 and D279, which mediate sodium and proton binding, could alter the pH-dependent activity of the antiporter. (E) G238, V240, D278 and R432, though they do not coordinate the sodium ion directly, are all located on the mobile and highly conserved core domain of the antiporter. (D) *Hs*NHA2 monomer, shown in cartoon representation, with position of various point mutations shown as yellow spheres. A sodium ion, observed in the experimental structure, is shown as a purple sphere and denotes the position of NHA2’s binding site.

The N-terminus of *Hs*NHA2 is poorly conserved except for the region between amino acids 45 and 60 (Fig. 7A). We identified two glutamates (E47, E56) and two lysines (K57, K58) within this region as important residues for *Hs*NHA2 activity (Fig. 7B), but not essential for the protein’s stability (Fig. 8D) or its localization in the plasma membrane (Fig. 7C). In particular, the replacement of these residues with an amino acid with the opposite charge resulted in *Hs*NHA2 with a decreased capacity to transport both its substrates, Li^+^ and Na^+^ (Fig. 7B).

In bison NHA2ΔN1-69, the TMS1 (aa 86 – 106) formed a unique homodimer interface with a large intracellular gap between the protomers, which closes in the presence of phosphoinositol lipids, and the lipid-compacted form was more active (Matsuoka et al., 2022). Our results confirm that truncations larger than 75 amino acids in *Hs*NHA2 in the immediate proximity of TMS1 (aa 82 – 105; Fig. 7A) led to the loss of function of the antiporter (Fig. 6A), most likely due to protein misfolding, and consequently mislocalization (Fig. 6C). The N-terminus seems to be also important for the stability of the protein, because with truncations larger than 70 amino acids we observed a decrease in the protein amount in cells (Fig. 6D). Nevertheless, our results clearly show that the N-terminal hydrophilic extension preceding TMS1 is an important part of the protein involved in the regulation of *Hs*NHA2’s function. Whether the structure of the conserved region between amino acids 45 – 60 or some posttranslational modification in this region are crucial for the regulation of *Hs*NHA2 remains to be investigated. Charged residues present in this segment could determine the flexibility of the N-terminus or they could be involved in intramolecular interactions and/or substrate attraction.

The core domain of Na^+^/H^+^ antiporters encapsulating the ion binding site includes two unwound transmembrane helices that cross each other in the middle of the membrane near the ion-binding site, creating an X-shaped structure, characteristic of the NhaA fold (Padan, 2014). As for the membrane domain of *Hs*NHA2, a comprehensive evolutionary analysis of 6537 representative CPAs revealed a sequence motif of 8 amino acids residing on different transmembrane segments, but spatially close to each other around the substrate binding site (Masrati et al., 2018). In *Hs*NHA2, this motif A2431V2442S2453P2464 …**-**5…D2786D2797….R4328 lacks the glutamate at position 5 present in other electrogenic CPA2 members. It was suggested that the electroneutrality of mammalian NHA2 is instead ensured by another glutamate (E215 in *Hs*NHA2) in TMS 5 (Fig. S1 and 8B) which forms a salt bridge with R432 at position 8 of the motif in TMS 12 (Masrati et al., 2018). The glutamate residue located in TMS 5 (Fig. 1, S1) is unique in the *Hs*NHA2 and its homologues and cannot be found in other CPA2 members (Masrati et al., 2018). CryoEM structures of bison and human NHA2ΔN proteins confirmed these two charged residues to be spatially close to each other (Fig. 8B; Protein Data Bank 7B4M/L; (Matsuoka et al., 2022)). Our mutagenesis studies of full version of *Hs*NHA2 corroborate the results obtained for bison NHA2ΔN that, in contrast to non-functional single-mutated versions (E215R or R432E), swapping up of E215 and R432 results in a Li^+^-selective transporter that transports Li^+^ within the pH range 4.0 – 6.0 (Fig. 4A, D). In addition, using fluorescence microscopy and immunodetection (Fig. 4B, C), we show that the interaction between these two residues is also important for its structural integrity, and hence for the trafficking of the protein.

Another residue that our study found to be highly important for the determination of *Hs*NHA2 transport properties is Pro246, located at position 4 in the conserved CPA motif close to the cation binding site D279 (Fig. 8C). The presence of proline in any TMS perturbs the α-helical structure due to the flexibility of the ring structure of its side chain and the elimination of helix backbone hydrogen bonds for the carbonyls at positions i-3 and i-4 (Visiers et al., 2000). In *Hs*NHA2, Pro246 is located in TMS 6 (Fig. 1, S1), which is one of the two unwound transmembrane segments that create the X-shaped structure in CPAs’ core domain (Fig. 8C). Depending on the nature of the amino acid substituted at this position, we observed changes in the substrate specificity (ability to recognize and transport Li^+^ or Na^+^) of the antiporter (Fig. 2A). Mutations of the corresponding proline residue to polar amino acids (Ser, Thr) in yeast Na^+^/H^+^ antiporters also resulted in altered substrate specificity towards larger cations (Kinclova-Zimmermannova et al., 2005). In human NHE1, the replacement of this residue with alanine or glycine resulted in the loss of transport activity (Counillon et al., 1997; Slepkov et al., 2005). In *Schizosacchromyces pombe* sod2 antiporter, mutations of corresponding proline affected its structure and localization (Ndayizeye et al., 2009). According to our results, the replacement of P246 with G, A, S or T did not change the targeting of *Hs*NHA2 to the plasma membrane or its stability (Fig. 2B, C). Nevertheless, in particular the presence of polar branched threonine resulted in an antiporter that transports better Na^+^ than Li^+^ and with altered capacity to transport protons, as it was highly active within the pH range 4.0 – 7.4, whilst the activity of the native antiporter could only be detected at pH 4.0 – 5.5 (Fig. 2A, B). Interestingly, measurements of Li^+^ binding by the purified *E. coli* NhaA showed that a variant mutated at the corresponding residue at position 4 of the conserved CPA motif to threonine (I134T) also exhibited a decrease in pH dependence, but the affinity of this mutated version to Li^+^ increased (Mondal et al., 2021). Here we show *in vivo* that proline 246 is involved in cation and proton coordination and binding in *Hs*NHA2.

Mutations of proline 246 to G, A, S or T provided the antiporter with highly increased resistance to the *Hs*NHA2-specific inhibitor phloretin (Fig. 3). It was found to efficiently inhibit e.g. the enzymatic activity of tyrosinase (J. Chen et al., 2020) or glucose transporters GLUT (Lin et al., 2016; L. Liu et al., 2022), and thus it influences various physiological processes (Casado-Diaz et al., 2021; Vadavanath Prabhakaran and Kozhiparambil Gopalan, 2021; Wu et al., 2018). Molecular docking showed that hydrophobic interaction and hydrogen bonding are the main non-covalent interaction forces between phloretin and tyrosinase (J. Chen et al., 2020). So far, the molecular mechanism of the phloretin inhibition of *Hs*NHA2 has not been studied. Here we show that the inhibition effect of phloretin did not change with mutations E215R + R432E in *Hs*NHA2, however it decreased significantly with alterations at the position of P246 (Fig. 3). Thus, our results indicate that phloretin acts directly in the core domain of *Hs*NHA2 and mutations of P246 possibly cause inhibitor resistance by altering TMS contacts near the substrate/inhibitor-binding pocket.

There are significant differences between the genomes of human individuals due to single-nucleotide variants (polymorphisms, SNPs). In this study we characterized the effect of six mutations existing as SNPs in human NHA2 (Dewey et al., 2016) on the antiporter’s transport activity, substrate specificity and localization (Fig. 5). The positions of mutated residues within the *Hs*NHA2 monomer in relation to the cation-binding site are shown in Fig. 8D. Residues G238, V240, D278 and R432, though they do not coordinate the transported cation (Na^+^ or Li^+^) directly, are all located on the mobile and highly conserved core domain of the antiporter (Fig. 8E). Our data show that this *in vivo* experimental approach can be highly useful for a rapid screening of SNP’s effect on *Hs*NHA2 activity. Here, it enabled us to distinguish the various effects of mutations on *Hs*NHA2 properties at the molecular level (e.g. effect on substrate specificity - R432Q; transport mechanism - D278G or protein folding and trafficking − G238R; Fig. 5). The mutation K382E was found to be associated with hypertension (Dewey et al., 2016). Positively charged residues are more abundant in the cytoplasmic side of many membrane proteins (“positive-inside rule”) and the substitution of such residue with an opposite charge could potentially affect the proper insertion and orientation of the protein to the membrane (Baker et al., 2017; Sipos and von Heijne, 1993; von Heijne, 2006). Notably, the positively charged K382 is located far from the cation-binding site, facing the cytoplasm (Fig. 8D), its replacement with negatively charged glutamate did not change the plasma-membrane localization of the antiporter (Fig. 5B), but it resulted in restricted substrate specificity for Li^+^ alone and very low transport activity (ability to improve the LiCl tolerance of cells) (Fig. 5A). On the other hand, the mutation V240L in the antiporter’s core domain (Fig. 8D, E) had only minor effect on *Hs*NHA2’s functionality (Fig. 5), probably, due to the conservative nature of this mutation. Furthermore, these results are in line with V240 positioning roughly 12 Å away from where the sodium ion binds (based on the Na^+^-bound structure of the *Hs*NHA2; PDB entry 7B4L; Fig. 8D). Thus, the combination of our experimental expression model for *Hs*NHA2 with molecular modelling can be used to characterize various pathology-related SNPs that exist in the *Hs*NHA2 gene in the future.

In general, our work provides new and important knowledge on using *S. cerevisiae* as a host for studying human membrane transporters. Specifically, it provides new valuable knowledge on the structure and function of the human Na^+^/H^+^ antiporter NHA2 that has broad physiological functions in the human body, and whose mal-functioning results in a growing list of pathologies. We identified several amino-acid residues important for *Hs*NHA2 substrate specificity, transport activity, stability and trafficking that had not been characterized previously. Furthermore, our optimized expression system of *Hs*NHA2 in yeast cells may be a powerful tool for the efficient screening of compounds that can activate/inhibit *Hs*NHA2 activity, for the selection of other inhibitor-resistant mutants to elucidate structure-activity relationships, or for discovering the molecular mechanism of inhibitors’ actions in *Hs*NHA2. Newly identified mutated versions of *Hs*NHA2 with higher activity and higher resistance to phloretin than the native antiporter may also serve as controls in future *Hs*NHA2 studies. Moreover, our findings concerning a unique regulatory/autoinhibitory role of the hydrophilic N-terminal part of the protein will definitely be useful for future studies of NHA2 regulation in human cells.

## Material and Methods

### Yeast strains and growth media

Alkali-metal-cation-sensitive *S. cerevisiae* W303-1A derivatives - BW31 (*ena1Δ*::*HIS3*::*ena4Δ nha1Δ*::*LEU2*) (Kinclova-Zimmermannova et al., 2005) or AB11c (*ena1Δ*::*HIS3*::*ena4Δ nha1Δ*::*LEU2 nhx1Δ::TRP1*) (Maresova and Sychrova, 2005) lacking genes encoding plasma-membrane Ena ATPases, plasma-membrane antiporter Nha1 or intracellular antiporter Nhx1 were used to characterize *Hs*NHA2 activity *in vivo*. Yeast cultures were routinely grown in YPD (Formedium, Hunstanton, UK) or YNB media (Difco, Sparks, MD, USA) at 30°C. YNB-based media (referred in this work as YNB-Pro) contained 0.17% YNB without amino acids and ammonium sulphate, 0.1% proline, and 2% or 4% glucose. Proline was used instead of ammonium sulphate as a nitrogen source. A mixture of appropriate auxotrophic supplements (adenine, tryptophan, leucine and histidine – each at a final concentration of 20 or 40 µg/ml) was added after autoclaving. YPD media were supplemented with extra adenine (final concentration 20 µg/ml) to avoid additional spontaneous mutagenesis caused by the *ade2* mutation present in both strains. Solid media were prepared by adding 2 or 3% (w/v) agar.

### Plasmids

The plasmids used in this study are listed in Table 1. All new plasmids were constructed by homologous recombination in the *S. cerevisiae* BW31 strain by using a series of overlapping PCR products and a linearized plasmid (pNHA1-985, pENA1 or pENA1-GFP; Table 1), in which the original gene was replaced by the corresponding *Hs*NHA2 cDNA or its appropriately truncated versions. The oligonucleotides used for PCR and the construction of particular plasmids are listed in Table S1. *Hs*NHA2 cDNA encoding full or N-terminally truncated versions of the antiporter were amplified from pTEF1-*Hs*NHA2-HA (Table 1). The sequence corresponding to GFP or the *ScTPS1* terminator were amplified from pGRU1 or YEp352-ZrSTL1-TPS1t-kanMX, respectively (Table 1). In all new plasmids, the expression of *Hs*NHA2 variants was under the control of the weak and constitutive *S. cerevisiae NHA1* promoter (Kinclova et al., 2001). The resulting plasmids were based on multicopy vectors YEp352 or pGRU1 (Table 1). Plasmids containing N-terminally GFP-tagged versions of *Hs*NHA2 (native or truncated) were constructed by inserting a GFP-encoding sequence in frame at the 5’-end. In p*Hs*NHA2-GFP, the GFP was added in frame at the 3’-end in front of the STOP codon. In p*Hs*NHA2*t* and pGFP-*Hs*NHA2*t* plasmids, the *ScTPS1* terminator was inserted behind the *Hs*NHA2 cDNA. To generate centromeric versions of p*Hs*NHA2*t* and pGFP-*Hs*NHA2*t*, the 2µ sequence was replaced by homologous recombination with the sequence corresponding to a centromere amplified from pRS316 (Sikorski and Hieter, 1989), and the resulting plasmids were named pC-*Hs*NHA2*t* and pC-GFP-*Hs*NHA2*t* (Table 1). *Escherichia coli* XL1-Blue (Agilent Technologies, Santa Clara, CA, USA) was used for pDNA amplification. Successful cloning was verified by restriction analysis and sequencing.

Point mutations were introduced into p*Hs*NHA2*t* or pGFP-p*Hs*NHA2*t* using a QuikChange XL Site-Directed Mutagenesis kit (Agilent Technologies) and the corresponding oligonucleotides (Table S1). The accuracy of mutations was confirmed by sequencing.

### Growth assays

To estimate the cell tolerance to different alkali-metal cations, YNB-Pro media were supplemented with LiCl, NaCl or KCl at the indicated concentration. HCl or TEA (triethylamine) were used to adjust the pH. Media with buffered pH were supplemented either with 20 mM MES (pH 5.5 and pH 6.0) or 20 mM MOPS (pH 7.0 and pH 7.4), and the required pH was adjusted as above. To test the activity inhibition of *Hs*NHA2 and its variants, media were supplemented with a solution of phloretin (Merck, Darmstadt, Germany; cat. no. P7912) dissolved in DMSO (100 mM stock solution). Tenfold serial dilutions of fresh cell suspensions (OD_600_= 2, Eppendorf BioPhotometer, Hamburg, Germany) were spotted on plates as indicated, and growth was monitored for 2-10 days. Representative results of at least three independent experiments are shown.

### Fluorescence microscopy

Microscopic images of yeast cells were acquired with an Olympus BX53 microscope with an Olympus DP73 camera (Olympus, Tokio, Japan). A Cool LED light source with 460 nm excitation and 515 nm emission was used to visualize *Hs*NHA2 and its variants or *Sc*Nha1 tagged with GFP (expressed from plasmids pGFP-*Hs*NHA2*t*, p*Hs*NHA2-GFP or pNHA1-985GFP, respectively, Table 1). Cells containing the corresponding multi-copy plasmids were grown overnight in YNB-Pro with 4% glucose and supplemented with adenine and tryptophan (final concentration 40 µg/ml), and leucine and histidine (final concentration 20 µg/ml). Cells were observed when they reached the exponential phase (OD_600_ ≈ 0.3 – 0.5). A Nomarski prism was used for whole-cell images.

### Cation loss determination

Sodium efflux was determined similarly as previously described (Kinclova-Zimmermannova et al., 2005). Cells were grown in YNB-Pro medium to the early exponential phase (OD_600_ ≈ 0.2-0.3, Spekol, Carl Zeiss, Oberkochen, Germany), harvested, resuspended and incubated in YNB-Pro supplemented with 100 mM NaCl at pH 7.0 (to preload them with sodium) for 60 minutes at 30°C. Cells were collected by centrifugation, washed with deionized water, and resuspended in an incubation buffer (10 mM Tris, 0.1 mM MgCl_2_, 2% glucose). The pH of the buffer was first brought down to 3.9 with citric acid and then adjusted to 4.0 with Ca(OH)_2_. The buffer was supplemented with 10 mM KCl to prevent Na^+^ reuptake. Samples of cell suspensions were withdrawn at intervals over 90 minutes. Cells were harvested by filtration (Millipore membrane filters, Merck-Millipore Co., Cork, Ireland), washed with 20 mM MgCl_2_, acid extracted, and the cation content in samples was determined by atomic absorption spectrometry (Kinclova-Zimmermannova et al., 2005). Data obtained in at least three independent experiments are shown as the average amount of sodium lost from cells over 90 minutes ± SD.

### Protein extraction and immunoblotting

Yeast cells containing the pGFP-*Hs*NHA2*t* plasmid or its variants were grown in liquid YNB-Pro media to the exponential phase (OD_600_ ≈ 0.6-0.8, Spekol, Carl Zeiss), collected by centrifugation, and washed once with deionized cold water. Cell pellets were kept at -80°C. Total extracts of proteins from cells were then prepared as described previously (Horak and Wolf, 2001). The RCDC protein assay (Bio-Rad, Hercules, CA, USA) was used for the quantification of proteins. Samples of total protein extracts (120 µg per sample) were separated by 10% SDS-PAGE and transferred to nitrocellulose membranes (Trans-Blot Turbo 0.2 µm nitrocellulose, Bio-Rad) using a Trans-Blot Turbo Transfer System (Bio-Rad). Membranes were incubated overnight at 4°C with an anti-GFP monoclonal antibody (Roche, Basel, Switzerland; dilution 1:500), washed and then incubated with a secondary anti-mouse IgG antibody tagged with a horseradish peroxidase (GE Healthcare, Chicago, IL, USA; dilution 1:10 000). Immunoreactive proteins were visualized with a Clarity Max Western ECL substrate kit (Bio-Rad) in ChemiDoc visualizer (Bio-Rad). To detect Pgk1 protein (as a protein loading control), the same membranes used for GFP-*Hs*NHA2 were reprobed after being incubated with an anti-Pgk1 mouse monoclonal antibody (Abcam, Cambridge, UK; dilution 1:20 000) for 1 hour at room temperature, incubated with the secondary antibody, and detected as above.

### *Hs*NHA2 evolutionary conservation analysis

The conservation analysis for *Hs*NHA2’s N-terminus was conducted using the ConSurf web server (http://dx.doi.org/10.1093/nar/gkw408). A total of 99 unique sequences were collected by HMMER from the UniRef90 database with sequence identity ranging between 35% to 95% and an E-value lower than 10^-5^.

### Statistics

Data presented are either from a representative experiment or from the mean of replicative values plus or minus standard deviation. Statistically significant differences were analysed by unpaired Student t-test using Microsoft Office Excel 2016 (** *p* < 0.01).

## Author contributions

Conceived and Designed Experiments: OZ, DV, and HS; Performed the experiments: DV, OZ, GM, VP. Analysed the Data: DV, OZ, GM. Drafted the Article: DV, OZ; Writing—review and editing: OZ, DV, HS, GM, and NB-T; Prepared the Digital Images: DV and GM.

## Acknowledgment

The technical assistance of Pavla Herynková is highly acknowledged. We thank Daniel Fuster, MD, for providing us the plasmid pTEF1-*Hs*NHA2-HA. We wish to thank Dr. Raghu Metpally and Prof. Rajini Rao for providing information about mutations identified as SNPs in *Hs*NHA2 that we characterized in this study. Prof. Rajini Rao and Prof. Pierre Falson are highly acknowledged also for a stimulating and fruitful discussion.

## Additional files

**Supplementary Table S1**: Oligonucleotides used in this study.

**Supplementary Figure S1**: The membrane topology of *Hs*NHA2.

**Supplementary Figure S2**: Comparison of localization of GFP-*Hs*NHA2 in two yeast backgrounds.

**Supplementary Figure S3**: N-terminal GFP-tagging does not change substrate specificity, but increases activity of *Hs*NHA2 mutated versions.

**Supplementary Figure S4**: N-terminal GFP-tagging does not change substrate specificity of **Hs**NHA2 versions with mutations that belong to known human SNPs.

**Supplementary Figure S5**: N-terminal GFP-tagging altered LiCl tolerance provided by *Hs*NHA2 versions truncated at N-terminus.

## References

Anderegg, M. A., Albano, G., Hanke, D., Deisl, C., Uehlinger, D. E., Brandt, S., Bhardwaj, R., Hediger, M. A., and Fuster, D. G. (2021). The sodium/proton exchanger NHA2 regulates blood pressure through a WNK4-NCC dependent pathway in the kidney. Kidney Int, 99(2), 350–363. doi:10.1016/j.kint.2020.08.023

Baker, J. A., Wong, W. C., Eisenhaber, B., Warwicker, J., and Eisenhaber, F. (2017). Erratum to: Charged residues next to transmembrane regions revisited: “Positive-inside rule” is complemented by the “negative inside depletion/outside enrichment rule”. BMC Biol, 15(1), 72. doi:10.1186/s12915-017-0410-6

Banuelos, M. A., Ruiz, M. C., Jimenez, A., Souciet, J. L., Potier, S., and Ramos, J. (2002). Role of the Nha1 antiporter in regulating K^+^ influx in *Saccharomyces cerevisiae*. Yeast, 19(1), 9–15. doi:10.1002/yea.799

Battaglino, R. A., Pham, L., Morse, L. R., Vokes, M., Sharma, A., Odgren, P. R., Yang, M., Sasaki, H., and Stashenko, P. (2008). NHA-oc/NHA2: A mitochondrial cation-proton antiporter selectively expressed in osteoclasts. Bone, 42(1), 180–192. doi:10.1016/j.bone.2007.09.046

Behzad, S., Sureda, A., Barreca, D., Nabavi, S. F., Rastrelli, L., and Nabavi, S. M. (2017). Health effects of phloretin: From chemistry to medicine. Phytochemistry Reviews, 16(3), 527–533. doi:10.1007/s11101-017-9500-x

Cagnac, O., Baghour, M., Jaime-Perez, N., Aranda-Sicilia, M. N., Sanchez-Romero, M. E., Rodriguez-Rosales, M. P., and Venema, K. (2020). Deletion of the N-terminal domain of the yeast vacuolar Na^+^, K^+^/H^+^ antiporter Vnx1p improves salt tolerance in yeast and transgenic arabidopsis. Yeast, 37(1), 173–185. doi:10.1002/yea.3450

Casado-Diaz, A., Rodriguez-Ramos, A., Torrecillas-Baena, B., Dorado, G., Quesada-Gomez, J. M., and Galvez-Moreno, M. A. (2021). Flavonoid phloretin inhibits adipogenesis and increases OPG expression in adipocytes derived from human bone-marrow mesenchymal stromal-cells. Nutrients, 13(11). doi:10.3390/nu13114185

Counillon, L., Noel, J., Reithmeier, R. A., and Pouyssegur, J. (1997). Random mutagenesis reveals a novel site involved in inhibitor interaction within the fourth transmembrane segment of the Na^+^/H^+^ exchanger-1. Biochemistry, 36(10), 2951–2959. doi:10.1021/bi9615405

Deisl, C., Anderegg, M., Albano, G., Luscher, B. P., Cerny, D., Soria, R., Bouillet, E., Rimoldi, S., Scherrer, U., and Fuster, D. G. (2016). Loss of sodium/hydrogen exchanger NHA2 exacerbates obesity- and aging-induced glucose intolerance in mice. PLoS One, 11(9), e0163568. doi:10.1371/journal.pone.0163568

Deisl, C., Simonin, A., Anderegg, M., Albano, G., Kovacs, G., Ackermann, D., Moch, H., Dolci, W., Thorens, B., M, A.H. and Fuster, D.G.. (2013). Sodium/hydrogen exchanger NHA2 is critical for insulin secretion in beta-cells. Proc Natl Acad Sci U S A, 110(24), 10004–10009. doi:10.1073/pnas.1220009110

Dewey, F. E., Murray, M. F., Overton, J. D., Habegger, L., Leader, J. B., Fetterolf, S. N., O’Dushlaine, C., Van Hout, C. V., Staples, J., Gonzaga-Jauregui, C., Metpally, R., Pendergrass, S. A., Giovanni, M. A., Kirchner, H. L., Balasubramanian, S., Abul-Husn, N. S., Hartzel, D. N., Lavage, D. R., Kost, K. A., Packer, J. S., Lopez, A. E., Penn, J., Mukherjee, S., Gosalia, N., Kanagaraj, M., Li, A. H., Mitnaul, L. J., Adams, J., Person, T. N., Praveen, K., Marcketta, A., Lebo, M. S., Austin-Tse, C. A., Mason-Suares, H. M., Bruse, S., Mellis, S., Phillips, R., Stahl, N., Murphy, A., Economides, A., Skelding, K. A., Still, C. D., Elmore, J. R., Borecki, I. B., Yancopoulos, G. D., Davis, F. D., Faucett, W. A., Gottesman, O., Ritchie, D., Shuldiner, A. R., Reid, J. G., Ledbetter, D. H., Baras, A., and Carey, D. J. (2016). Distribution and clinical impact of functional variants in 50,726 whole-exome sequences from the discovehr study. Science, 354(6319). doi:10.1126/science.aaf6814

Dong, Y., Gao, Y., Ilie, A., Kim, D., Boucher, A., Li, B., Zhang, X. C., Orlowski, J., and Zhao, Y. (2021). Structure and mechanism of the human NHE1-CHP1 complex. Nat Commun, 12(1), 3474. doi:10.1038/s41467-021-23496-z

Duskova, M., Ferreira, C., Lucas, C., and Sychrova, H. (2015). Two glycerol uptake systems contribute to the high osmotolerance of *Zygosaccharomyces rouxii*. Mol Microbiol, 97(3), 541–559. doi:10.1111/mmi.13048

Flegelova, H., Haguenauer-Tsapis, R., and Sychrova, H. (2006). Heterologous expression of mammalian Na^+^/H^+^ antiporters in *Saccharomyces cerevisiae*. Biochim Biophys Acta, 1760(3), 504–516. doi:10.1016/j.bbagen.2006.01.014

Flegelova, H., and Sychrova, H. (2005). Mammalian NHE2 Na^+^/H^+^ exchanger mediates efflux of potassium upon heterologous expression in yeast. FEBS Lett, 579(21), 4733–4738. doi:10.1016/j.febslet.2005.07.046

Fuster, D. G., Zhang, J., Shi, M., Bobulescu, I. A., Andersson, S., and Moe, O. W. (2008). Characterization of the sodium/hydrogen exchanger NHA2. J Am Soc Nephrol, 19(8), 1547–1556. doi:10.1681/ASN.2007111245

Goswami, P., Paulino, C., Hizlan, D., Vonck, J., Yildiz, O., and Kuhlbrandt, W. (2011). Structure of the archaeal Na^+^/H^+^ antiporter NhaP1 and functional role of transmembrane helix 1. Embo J, 30(2), 439–449. doi:10.1038/emboj.2010.321

Ha, B. G., Hong, J. M., Park, J. Y., Ha, M. H., Kim, T. H., Cho, J. Y., Ryoo, H. M., Choi, J. Y., Shin, H. I., Chun, S. Y., Kim, S. Y., and Park, E. K. (2008). Proteomic profile of osteoclast membrane proteins: Identification of Na^+^/H^+^ exchanger domain containing 2 and its role in osteoclast fusion. Proteomics, 8(13), 2625–2639. doi:10.1002/pmic.200701192

Hendus-Altenburger, R., Kragelund, B. B., and Pedersen, S. F. (2014). Structural dynamics and regulation of the mammalian SLC9A family of Na^+^/H^+^ exchangers. Exchangers, 73, 69–148. doi:10.1016/B978-0-12-800223-0.00002-5

Hill, J. E., Myers, A. M., Koerner, T. J., and Tzagoloff, A. (1986). Yeast/*E. coli* shuttle vectors with multiple unique restriction sites. Yeast, 2, 163–167.

Hofstetter, W., Siegrist, M., Simonin, A., Bonny, O., and Fuster, D. G. (2010). Sodium/hydrogen exchanger NHA2 in osteoclasts: Subcellular localization and role *in vitro* and *in vivo*. Bone, 47(2), 331–340. doi:10.1016/j.bone.2010.04.605

Horak, J., and Wolf, D. H. (2001). Glucose-induced monoubiquitination of the *Saccharomyces cerevisiae* galactose transporter is sufficient to signal its internalization. Journal of Bacteriology, 183(10), 3083–3088. doi:10.1128/JB.183.10.3083-3088.2001

Hunte, C., Screpanti, E., Venturi, M., Rimon, A., Padan, E., and Michel, H. (2005). Structure of a Na^+^/H^+^ antiporter and insights into mechanism of action and regulation by pH. Nature, 435(7046), 1197–1202. doi:10.1038/nature03692

Charles, J. F., Coury, F., Sulyanto, R., Sitara, D., Wu, J., Brady, N., Tsang, K., Sigrist, K., Tollefsen, D. M., He, L., Storm, D., and Aliprantis, A. O. (2012). The collection of NFATc1-dependent transcripts in the osteoclast includes numerous genes non-essential to physiologic bone resorption. Bone, 51(5), 902–912. doi:10.1016/j.bone.2012.08.113

Chen, J., Li, Q., Ye, Y., Huang, Z., Ruan, Z., and Jin, N. (2020). Phloretin as both a substrate and inhibitor of tyrosinase: Inhibitory activity and mechanism. Spectrochim Acta A Mol Biomol Spectrosc, 226, 117642. doi:10.1016/j.saa.2019.117642

Chen, S. R., Chen, M., Deng, S. L., Hao, X. X., Wang, X. X., and Liu, Y. X. (2016). Sodium-hydrogen exchanger NHA1 and NHA2 control sperm motility and male fertility. Cell Death Dis, 7, e2152. doi:10.1038/cddis.2016.65

Chintapalli, V. R., Kato, A., Henderson, L., Hirata, T., Woods, D. J., Overend, G., Davies, S. A., Romero, M. F., and Dow, J. A. (2015). Transport proteins NHA1 and NHA2 are essential for survival, but have distinct transport modalities. Proc Natl Acad Sci U S A, 112(37), 11720–11725. doi:10.1073/pnas.1508031112

Kinclova-Zimmermannova, O., Zavrel, M., and Sychrova, H. (2005). Identification of conserved prolyl residue important for transport activity and the substrate specificity range of yeast plasma membrane Na^+^/H^+^ antiporters. J Biol Chem, 280(34), 30638–30647. doi:10.1074/jbc.M506341200

Kinclova, O., Ramos, J., Potier, S., and Sychrova, H. (2001). Functional study of the *Saccharomyces cerevisiae* Nha1p C-terminus. Mol Microbiol, 40(3), 656–668. doi:10.1046/j.1365-2958.2001.02412.x

Kondapalli, K. C., Kallay, L. M., Muszelik, M., and Rao, R. (2012). Unconventional chemiosmotic coupling of NHA2, a mammalian Na^+^/H^+^ antiporter, to a plasma membrane H^+^ gradient. J Biol Chem, 287(43), 36239–36250. doi:10.1074/jbc.M112.403550

Kondapalli, K. C., Todd Alexander, R., Pluznick, J. L., and Rao, R. (2017). NHA2 is expressed in distal nephron and regulated by dietary sodium. J Physiol Biochem, 73(2), 199–205. doi:10.1007/s13105-016-0539-8

Lee, C., Kang, H. J., von Ballmoos, C., Newstead, S., Uzdavinys, P., Dotson, D. L., Iwata, S., Beckstein, O., Cameron, A. D., and Drew, D. (2013). A two-domain elevator mechanism for sodium/proton antiport. Nature, 501(7468), 573–577. doi:10.1038/nature12484

Lee, S. H., Kim, T., Park, E. S., Yang, S., Jeong, D., Choi, Y., and Rho, J. (2008). NHE10, an osteoclast-specific member of the Na^+^/H^+^ exchanger family, regulates osteoclast differentiation and survival [corrected]. Biochem Biophys Res Commun, 369(2), 320–326. doi:10.1016/j.bbrc.2008.01.168

Lin, S. T., Tu, S. H., Yang, P. S., Hsu, S. P., Lee, W. H., Ho, C. T., Wu, C. H., Lai, Y. H., Chen, M. Y., and Chen, L. C. (2016). Apple polyphenol phloretin inhibits colorectal cancer cell growth via inhibition of the type 2 glucose transporter and activation of p53-mediated signaling. J Agric Food Chem, 64(36), 6826–6837. doi:10.1021/acs.jafc.6b02861

Liu, H. M., He, J. Y., Zhang, Q., Lv, W. Q., Xia, X., Sun, C. Q., Zhang, W. D., and Deng, H. W. (2018). Improved detection of genetic loci in estimated glomerular filtration rate and type 2 diabetes using a pleiotropic cfdr method. Mol Genet Genomics, 293(1), 225–235. doi:10.1007/s00438-017-1381-6

Liu, L., Xie, H., Zhao, S., and Huang, X. (2022). The GLUT1-mtORC1 axis affects odontogenic differentiation of human dental pulp stem cells. Tissue Cell, 76, 101766 doi:10.1016/j.tice.2022.101766

Maresova, L., and Sychrova, H. (2005). Physiological characterization of *Saccharomyces cerevisiae kha1* deletion mutants. Mol Microbiol, 55(2), 588–600.

Masrati, G., Dwivedi, M., Rimon, A., Gluck-Margolin, Y., Kessel, A., Ashkenazy, H., Mayrose, I., Padan, E., and Ben-Tal, N. (2018). Broad phylogenetic analysis of cation/proton antiporters reveals transport determinants. Nat Commun, 9(1), 4205. doi:10.1038/s41467-018-06770-5

Matsuoka, R., Fudim, R., Jung, S., Zhang, C., Bazzone, A., Chatzikyriakidou, Y., Robinson, C. V., Nomura, N., Iwata, S., Landreh, M., Orellana, L., Beckstein, O., and Drew, D. (2022). Structure, mechanism and lipid-mediated remodeling of the mammalian Na^+^/H^+^ exchanger NHA2. Nat Struct Mol Biol, 29(2), 108–120. doi:10.1038/s41594-022-00738-2

Mondal, R., Rimon, A., Masrati, G., Ben-Tal, N., Friedler, A., and Padan, E. (2021). Towards molecular understanding of the pH dependence characterizing NhaA of which structural fold is shared by other transporters. J Mol Biol, 433(19), 167156. doi:10.1016/j.jmb.2021.167156

Ndayizeye, M., Touret, N., and Fliegel, L. (2009). Proline 146 is critical to the structure, function and targeting of sod2, the Na^+^/H^+^ exchanger of *Schizosaccharomyces pombe*. Biochim Biophys Acta, 1788(5), 983–992. doi:10.1016/j.bbamem.2009.01.001

Padan, E. (2014). Functional and structural dynamics of NhaA, a prototype for Na^+^ and H^+^ antiporters, which are responsible for Na^+^ and H^+^ homeostasis in cells. Biochim Biophys Acta, 1837(7), 1047–1062. doi:10.1016/j.bbabio.2013.12.007

Padan, E., and Landau, M. (2016). Sodium-proton (Na^+^/H^+^) antiporters: Properties and roles in health and disease. Met Ions Life Sci, 16, 391–458. doi:10.1007/978-3-319-21756-7_12

Paulino, C., Wohlert, D., Kapotova, E., Yildiz, O., and Kuhlbrandt, W. (2014). Structure and transport mechanism of the sodium/proton antiporter MjNhaP1. Elife, 3, e03583. doi:10.7554/eLife.03583

Pedersen, S. F., and Counillon, L. (2019). The SLC9A-C mammalian Na^+^/H^+^ exchanger family: Molecules, mechanisms, and physiology. Physiol Rev, 99(4), 2015–2113. doi:10.1152/physrev.00028.2018

Pittman, J. K., Sreevidya, C. S., Shigaki, T., Ueoka-Nakanishi, H., and Hirschi, K. D. (2002). Distinct N-terminal regulatory domains of Ca^2+^/H^+^ antiporters. Plant Physiol, 130(2), 1054–1062. doi:10.1104/pp.008193

Prasad, H., Dang, D. K., Kondapalli, K. C., Natarajan, N., Cebotaru, V., and Rao, R. (2019). NHA2 promotes cyst development in an *in vitro* model of polycystic kidney disease. J Physiol, 597(2), 499–519. doi:10.1113/JP276796

Schushan, M., Xiang, M., Bogomiakov, P., Padan, E., Rao, R., and Ben-Tal, N. (2010). Model-guided mutagenesis drives functional studies of human NHA2, implicated in hypertension. J Mol Biol, 396(5), 1181–1196. doi:10.1016/j.jmb.2009.12.055

Sikorski, R. S., and Hieter, P. (1989). A system of shuttle vectors and yeast host strains designed for efficient manipulation of DNA in *Saccharomyces cerevisiae*. Genetics, 122(1), 19–27.

Sipos, L., and von Heijne, G. (1993). Predicting the topology of eukaryotic membrane proteins. Eur J Biochem, 213(3), 1333–1340. doi:10.1111/j.1432-1033.1993.tb17885.x

Slepkov, E. R., Rainey, J. K., Li, X., Liu, Y., Cheng, F. J., Lindhout, D. A., Sykes, B. D., and Fliegel, L. (2005). Structural and functional characterization of transmembrane segment IV of the NHE1 isoform of the Na^+^/H^+^ exchanger. J Biol Chem. doi:10.1074/jbc.M409608200

Smidova, A., Stankova, K., Petrvalska, O., Lazar, J., Sychrova, H., Obsil, T., Zimmermannova, O., and Obsilova, V. (2019). The activity of *Saccharomyces cerevisiae* Na^+^, K^+^/H^+^ antiporter Nha1 is negatively regulated by 14-3-3 protein binding at serine 481. Biochim Biophys Acta Mol Cell Res, 1866(12), 118534. doi:10.1016/j.bbamcr.2019.118534

Stribny, J., Thines, L., Deschamps, A., Goffin, P., and Morsomme, P. (2020). The human golgi protein TMEM165 transports calcium and manganese in yeast and bacterial cells. J Biol Chem, 295(12), 3865–3874. doi:10.1074/jbc.RA119.012249

Su, P., Wu, H., Wang, M., Cai, L., Liu, Y., and Chen, L. M. (2021). IRBIT activates NBCe1-b by releasing the auto-inhibition module from the transmembrane domain. J Physiol, 599(4), 1151–1172. doi:10.1113/JP280578

Thines, L., Deschamps, A., Stribny, J., and Morsomme, P. (2019). Yeast as a tool for deeper understanding of human manganese-related diseases. Genes (Basel), 10(7). doi:10.3390/genes10070545

Uzdavinys, P., Coincon, M., Nji, E., Ndi, M., Winkelmann, I., von Ballmoos, C., and Drew, D. (2017). Dissecting the proton transport pathway in electrogenic Na^+^/H^+^ antiporters. Proc Natl Acad Sci U S A, 114(7), E1101–E1110. doi:10.1073/pnas.1614521114

Vadavanath Prabhakaran, V., and Kozhiparambil Gopalan, R. (2021). Phloretin alleviates arsenic trioxide-induced apoptosis of H9c2 cardiomyoblasts via downregulation in Ca^2+^/calcineurin/NFATc pathway and inflammatory cytokine release. Cardiovasc Toxicol, 21(8), 642–654. doi:10.1007/s12012-021-09655-0

Varadi, M., Anyango, S., Deshpande, M., Nair, S., Natassia, C., Yordanova, G., Yuan, D., Stroe, O., Wood, G., Laydon, A., Zidek, A., Green, T., Tunyasuvunakool, K., Petersen, S., Jumper, J., Clancy, E., Green, R., Vora, A., Lutfi, M., Figurnov, M., Cowie, A., Hobbs, N., Kohli, P., Kleywegt, G., Birney, E., Hassabis, D., and Velankar, S. (2022). Alphafold protein structure database: Massively expanding the structural coverage of protein-sequence space with high-accuracy models. Nucleic Acids Res, 50(D1), D439–D444. doi:10.1093/nar/gkab1061

Visiers, I., Braunheim, B. B., and Weinstein, H. (2000). Prokink: A protocol for numerical evaluation of helix distortions by proline. Protein Eng, 13(9), 603–606. doi:10.1093/protein/13.9.603

von Heijne, G. (2006). Membrane-protein topology. Nat Rev Mol Cell Biol, 7(12), 909–918. doi:10.1038/nrm2063

Winklemann, I., Matsuoka, R., Meier, P. F., Shutin, D., Zhang, C., Orellana, L., Sexton, R., Landreh, M., Robinson, C. V., Beckstein, O., and Drew, D. (2020). Structure and elevator mechanism of the mammalian sodium/proton exchanger NHE9. Embo J, 39(24), e105908. doi:10.15252/embj.2020105908

Wohlert, D., Kuhlbrandt, W., and Yildiz, O. (2014). Structure and substrate ion binding in the sodium/proton antiporter panhap. Elife, 3, e03579. doi:10.7554/eLife.03579

Wu, K. H., Ho, C. T., Chen, Z. F., Chen, L. C., Whang-Peng, J., Lin, T. N., and Ho, Y. S. (2018). The apple polyphenol phloretin inhibits breast cancer cell migration and proliferation via inhibition of signals by type 2 glucose transporter. J Food Drug Anal, 26(1), 221–231. doi:10.1016/j.jfda.2017.03.009

Xiang, M., Feng, M., Muend, S., and Rao, R. (2007). A human Na^+^/H^+^ antiporter sharing evolutionary origins with bacterial NhaA may be a candidate gene for essential hypertension. Proc Natl Acad Sci U S A, 104(47), 18677–18681. doi:10.1073/pnas.0707120104

Yamanishi, M., Katahira, S., and Matsuyama, T. (2011). *TPS1* terminator increases mRNA and protein yield in a S*accharomyces cerevisiae* expression system. Biosci Biotechnol Biochem, 75(11), 2234–2236. doi:10.1271/bbb.110246

Zhao, J., Hyman, L., and Moore, C. (1999). Formation of mRNA 3’ ends in eukaryotes: Mechanism, regulation, and interrelationships with other steps in mRNA synthesis. Microbiol Mol Biol Rev, 63(2), 405–445. doi:10.1128/MMBR.63.2.405-445.1999

