## Supplementary material for "Allosteric links between the hydrophilic N-terminus and transmembrane core of human Na^+^/H^+^ antiporter NHA2"

Diego Velázquez[1], Vojtěch Průša[1], Gal Masrati[2], Hana Sychrova[1], Nir Ben-Tal[2], Olga Zimmermannova[1*]

[1]Laboratory of Membrane Transport, Institute of Physiology of the Czech Academy of Sciences, Prague 4, Czech Republic

[2]Department of Biochemistry and Molecular Biology, George S. Wise Faculty of Life Sciences, Tel-Aviv University, Tel-Aviv, Israel

E-mail addresses to other authors:

Diego Vélazquez (D.V.):

Vojtěch Průša (V.P.):

Gal Masrati (G.M.):

Hana Sychrova (H.S.):

Nir Ben-Tal (N.B-T.):

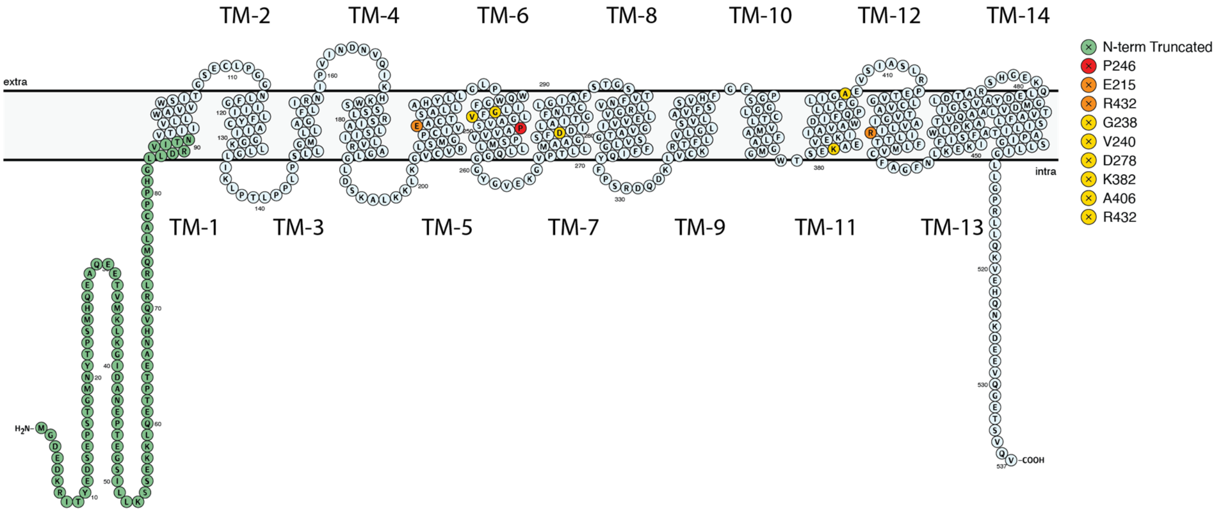


**Figure S1: The membrane topology of *Hs*NHA2.** *Hs*NHA2 orientation in the membrane is shown in 2-dimensional representation with the extracellular matrix at the top and the cytoplasm at the bottom. Transmembrane helices are numbered 1-to-14. The N-terminus region, where various truncations were made, is colored green. The position of P246, which is involved in pH regulation, is highlighted in red, E215 and R432 that form a conserved salt bridge are colored orange, and the positions of other human variants are highlighted in yellow.

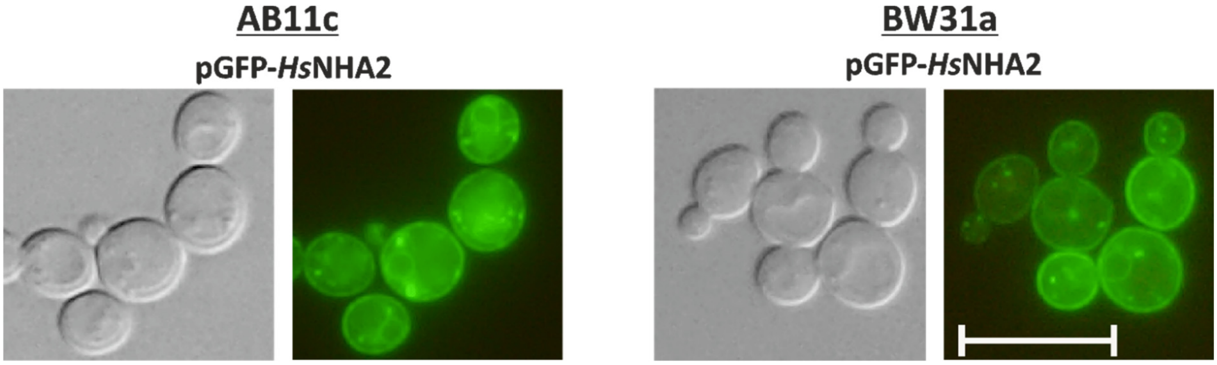


**Figure S2: Comparison of localization of GFP-*Hs*NHA2 in two yeast backgrounds.** Transformants of AB11c (*ena1-4Δ nha1Δ nhx1Δ*) or BW31 (*ena1-4Δ nha1Δ*) expressing GFP-*Hs*NHA2 from the pGFP-*Hs*NHA2 plasmid were grown in YNB-Pro (4% glucose) to the exponential phase and observed under a fluorescence microscope (right). A Nomarski prism was used for whole-cell imaging (left). The scale bar corresponds to 10 µm.

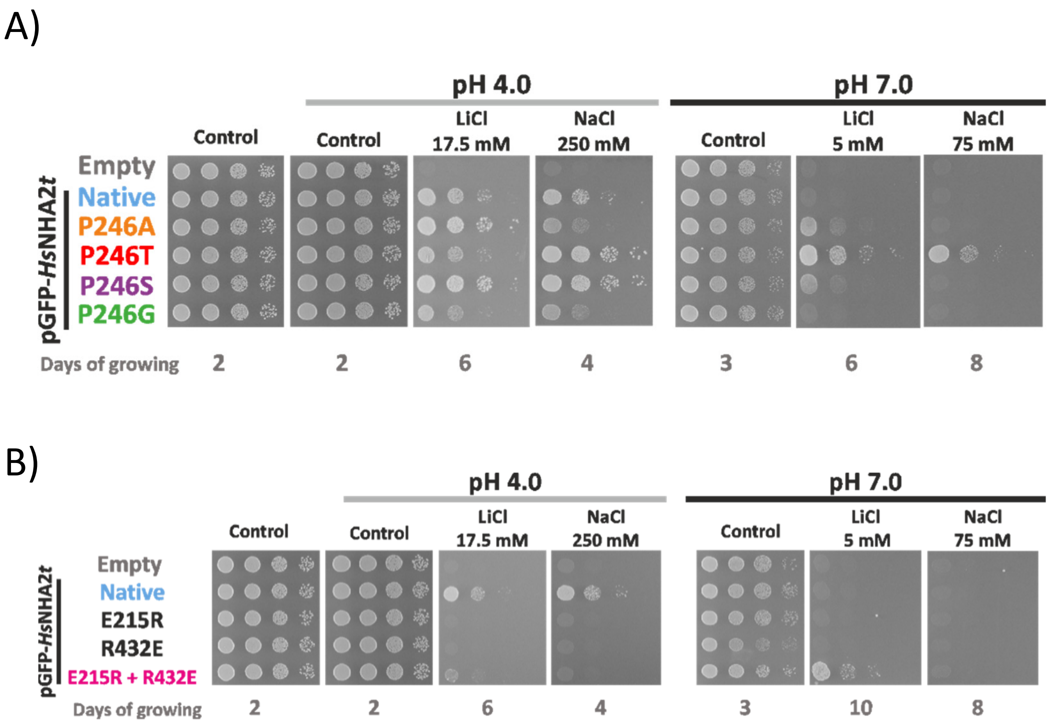


**Figure S3: N-terminal GFP-tagging does not change substrate specificity, but increases activity of *Hs*NHA2 mutated versions.** Salt tolerance of *S. cerevisiae* BW31 cells containing empty vector or expressing GFP-*Hs*NHA2 or one of four GFP-*Hs*NHA2 versions with single point mutations P246A, T, S or G (A) or versions E215R, R432E or E215R + R432E (B) from pGFP-*Hs*NHA2*t* plasmid. Cells were grown on non-buffered YNB-Pro plates with the pH adjusted to 4.0 or 7.0 and supplemented with LiCl or NaCl as indicated. Plates were incubated at 30 °C and photographed on the indicated day.

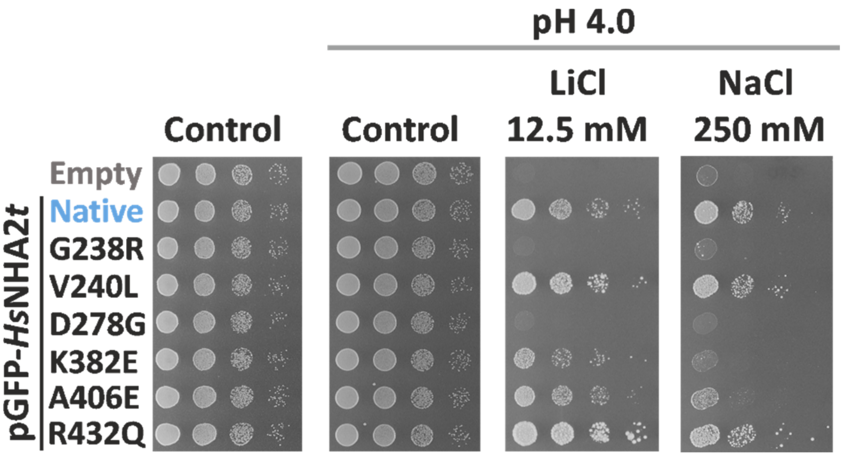


**Figure S4: N-terminal GFP-tagging does not change substrate specificity of *Hs*NHA2 versions with mutations that belong to known human SNPs.** The salt tolerance of *S. cerevisiae* BW31 cells containing the empty vector or expressing the native GFP-*Hs*NHA2 or one of six GFP-*Hs*NHA2 mutated versions - G238R, V240L, D278G, K382E, A406E and R432Q from the pGFP-*Hs*NHA2*t* plasmid. Cells were grown on non-buffered YNB-Pro plates with the pH adjusted to 4.0 and supplemented with LiCl or NaCl as indicated. Growth was monitored for 2 (control) or 7 (LiCl or NaCl) days at 30°C.

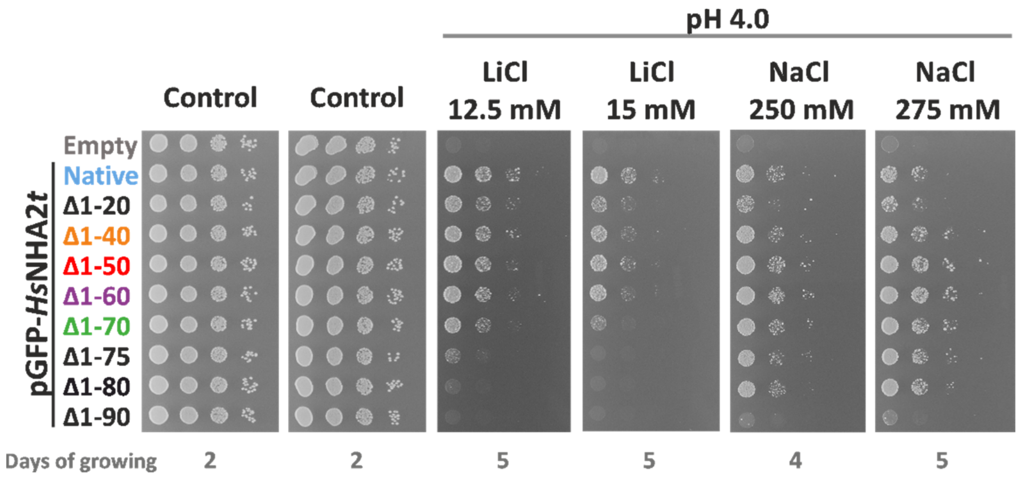


**Figure S5: N-terminal GFP-tagging altered LiCl tolerance provided by *Hs*NHA2 versions truncated at N-terminus.** The salt tolerance of *S. cerevisiae* BW31 cells containing the empty vector or expressing the tagged GFP-*Hs*NHA2 or N-terminal truncated GFP-*Hs*NHA2 versions from the pGFP-*Hs*NHA2*t*. Cells were grown on non-buffered YNB-Pro plates with the pH adjusted to 4.0 and supplemented with LiCl or NaCl as indicated. Plates were incubated at 30 °C and photographed on the indicated day.
